# Unique structural features govern the activity of a human mitochondrial AAA+ disaggregase, Skd3

**DOI:** 10.1101/2022.02.17.480866

**Authors:** Ryan R. Cupo, Alexandrea N. Rizo, Gabriel A. Braun, Eric Tse, Edward Chuang, Daniel R. Southworth, James Shorter

**Affiliations:** Department of Biochemistry and Biophysics, Perelman School of Medicine, University of Pennsylvania, Philadelphia, PA, U.S.A; Pharmacology Graduate Group, Perelman School of Medicine, University of Pennsylvania, Philadelphia, PA, U.S.A; Department of Biochemistry and Biophysics, Institute for Neurodegenerative Diseases, University of California, San Francisco, San Francisco, CA, U.S.A; Graduate Program in Chemical Biology, University of Michigan, Ann Arbor, MI, USA; Chemistry and Chemical Biology Graduate Program, University of California, San Francisco, CA, U.S.A

## Abstract

The AAA+ protein, Skd3 (human *CLPB*), solubilizes proteins in the mitochondrial intermembrane space, which is critical for human health. Skd3 variants with impaired protein-disaggregase activity cause severe congenital neutropenia (SCN) and 3-methylglutaconic aciduria type 7 (MGCA7). Yet how Skd3 disaggregates proteins remains poorly understood. Here, we report a high-resolution structure of a Skd3-substrate complex. Skd3 adopts a spiral hexameric arrangement that engages substrate via pore-loop interactions in the nucleotide-binding domain (NBD). Unexpectedly, substrate-bound Skd3 hexamers stack head-to-head via unique, adaptable ankyrin-repeat domain (ANK)-mediated interactions to form dodecamers. Deleting the ANK-linker region reduces dodecamerization and disaggregase activity. We elucidate apomorphic features of the Skd3 NBD and C-terminal domain that regulate disaggregase activity. We also define how Skd3 subunits collaborate to disaggregate proteins. Importantly, SCN-linked subunits sharply inhibit disaggregase activity, whereas MGCA7-linked subunits do not. Our findings illuminate Skd3 structure and mechanism, explain SCN and MGCA7 inheritance patterns, and suggest therapeutic strategies.

## Introduction

Protein aggregation and aberrant phase transitions can have deleterious consequences, including neurodegenerative disease (Darling and Shorter, 2021; Eisele et al., 2015). Thus, specialized protein disaggregases have evolved to safely reverse protein aggregation and restore resolubilized proteins to native structure and function (Fare and Shorter, 2021). These include ATP-independent systems, such as DAXX, TRIMs, and nuclear-import receptors, as well as ATP-dependent systems, including specific AAA+ (<u>A</u>TPases <u>a</u>ssociated with diverse cellular <u>a</u>ctivities) proteins such as Hsp104 and Skd3 (human *CLPB*) (Cupo and Shorter, 2020b; Guo et al., 2018; Huang et al., 2021; Shorter, 2017; Shorter and Southworth, 2019; Zhu et al., 2020).

AAA+ proteins couple ATP hydrolysis to mechanical work to power various energetically challenging tasks, including protein disaggregation (Puchades et al., 2020; Shorter and Southworth, 2019). For example, Hsp104, a hexameric, double AAA+ ring disaggregase found in all non-metazoan eukaryotes, can disassemble stable amyloids, prions, amorphous aggregates, toxic oligomers, and heat-induced condensates (DeSantis et al., 2012; Lo Bianco et al., 2008; Shorter and Lindquist, 2004, 2006; Sweeny et al., 2015; Yoo et al., 2022). To disaggregate proteins, Hsp104 and its bacterial homolog, ClpB, translocate polypeptides into their central channels using tyrosine-bearing pore loops that grip substrate (Gates et al., 2017; Rizo et al., 2019; Shorter and Southworth, 2019; Yokom et al., 2016). Curiously, despite having potent neuroprotective activity when expressed in animal models (Cushman-Nick et al., 2013; Lo Bianco et al., 2008), Hsp104 was lost during the evolutionary transition from protozoa to metazoa, as was its mitochondrial counterpart, Hsp78 (Erives and Fassler, 2015). However, humans express Skd3, a single AAA+ ring disaggregase found in the mitochondrial intermembrane space, which first appears in evolution alongside Hsp104 and Hsp78 in the closest extant protozoan relatives of animals (Erives and Fassler, 2015). Skd3 is related to Hsp104 and Hsp78 via its HCLR clade AAA+ domain, but otherwise shares limited homology (Erives and Fassler, 2015; Erzberger and Berger, 2006; Perier et al., 1995; Seraphim and Houry, 2020).

Skd3 functions to maintain protein solubility in the mitochondrial intermembrane space and ensures mitochondrial functionality (Chen et al., 2019; Cupo and Shorter, 2020b; Warren et al., 2022). Indeed, Skd3 exhibits potent protein-disaggregase activity and plays a critical role in human health (Cupo and Shorter, 2020b; Warren et al., 2022). Autosomal dominant mutations in Skd3 that impair disaggregase activity cause severe congenital neutropenia (SCN) (Warren et al., 2022). SCN is a rare bone marrow failure syndrome that presents with impaired neutrophil maturation (Skokowa et al., 2017). Due to low neutrophil counts, SCN patients are prone to life-threatening infections early in life and exhibit increased propensity for myelodysplastic syndromes or acute myeloid leukemia (Skokowa et al., 2017). By contrast, autosomal recessive or distinct biallelic mutations in Skd3 that impair disaggregase activity underlie 3-methylglutaconic aciduria type 7 (MGCA7) (Cupo and Shorter, 2020b; Kanabus et al., 2015; Kiykim et al., 2016; Pronicka et al., 2017; Saunders et al., 2015; Wortmann et al., 2016; Wortmann et al., 2021; Wortmann et al., 2015; Zhang et al., 2020). MGCA7 presents with elevated levels of 3-methylglutaconic acid, neurologic deterioration, and neutropenia (Wortmann et al., 2016; Wortmann et al., 2015). Patients present with infantile onset of a progressive encephalopathy with movement abnormalities and delayed psychomotor development, which can be accompanied by cataracts, seizures, and recurrent infections (Wortmann et al., 2016; Wortmann et al., 2015). In severe cases, afflicted infants die within a few weeks (Wortmann et al., 2016; Wortmann et al., 2015). There are no effective therapeutics for severe MGCA7. Finally, Skd3 has emerged as a therapeutic target to inhibit in prostate cancer and Venetoclax-resistant acute myeloid leukemia (Chen et al., 2019; Pudova et al., 2020).

Despite the importance of Skd3 disaggregase activity for human health, little is known about the mechanism of action or structure of Skd3 (Cupo and Shorter, 2020b). Skd3 harbors an N-terminal mitochondrial targeting signal, which is cleaved by mitochondrial processing peptidase (MPP) upon import into the mitochondria (Cupo and Shorter, 2020b; Wortmann et al., 2015). Skd3 then has a hydrophobic autoinhibitory peptide, which is removed by PARL, a rhomboid protease in the mitochondrial inner membrane (Saita et al., 2017). Removal of this peptide increases Skd3 disaggregase activity by more than 10-fold (Cupo and Shorter, 2020b). Thus, Skd3 is only fully activated upon reaching its final destination in the mitochondrial intermembrane space. After these processing events, the mature form of Skd3 contains an ankyrin-repeat domain (ANK), a nucleotide-binding domain (NBD) from the HCLR clade of the AAA+ family (Erzberger and Berger, 2006; Seraphim and Houry, 2020), and a short C-terminal domain (CTD; Figure 1A).

**Figure 1:**
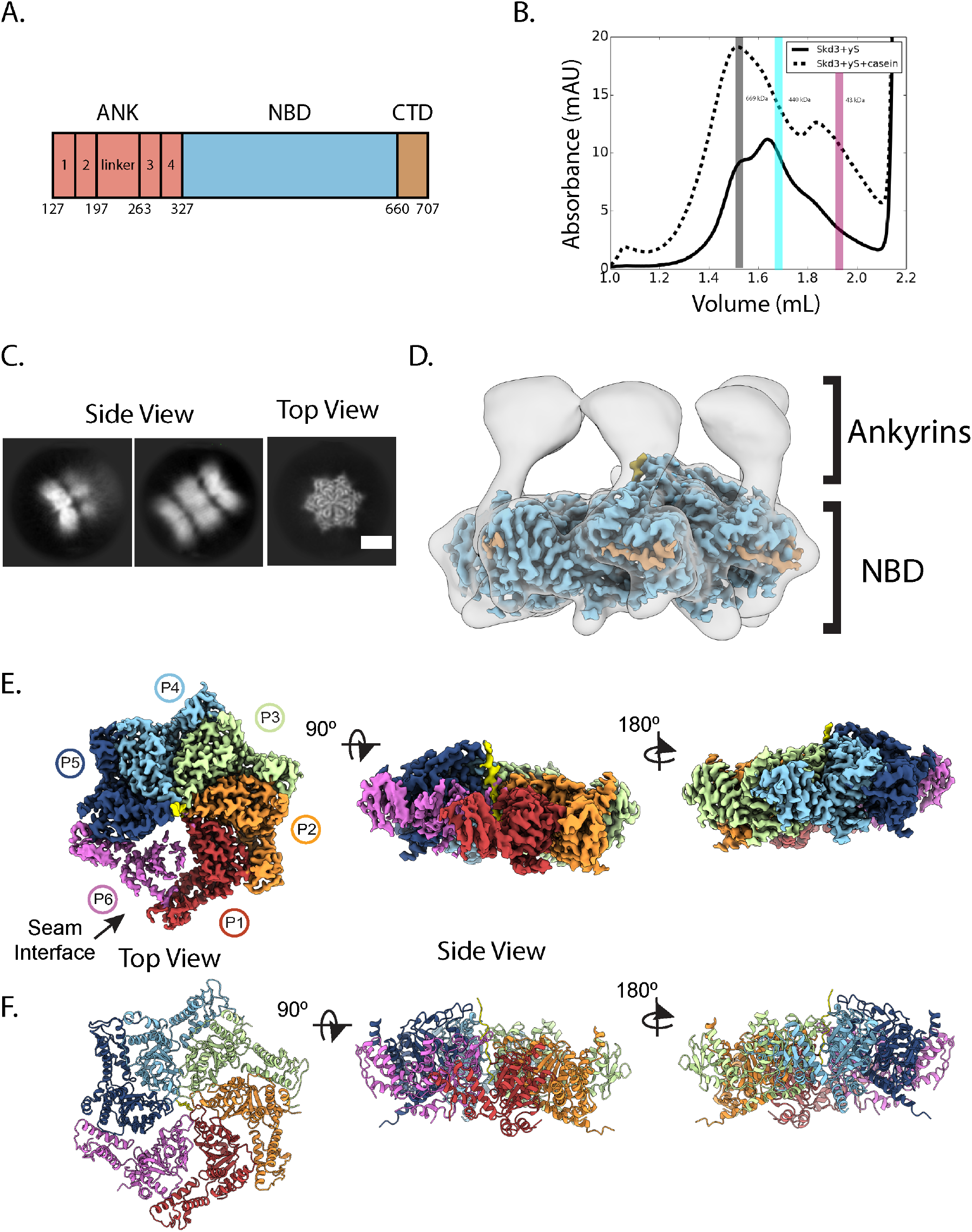
Structure of _PARL_Skd3. **(A)** Domain schematic of _PARL_Skd3. **(B)** SEC of _PARL_Skd3 incubated with ATPγS (solid) or ATPγS and FITC-casein (dashed). Vertical bars indicate elution position of molecular-weight standards: thyroglobulin (669 kDa), ferritin (440 kDa), and ovalbumin (43 kDa), and are representative of _PARL_Skd3 dodecamer (grey), hexamer (cyan), or monomer (magenta) size. **(C)** Representative cryo-EM 2D class averages of the _PARL_Skd3:casein:ATPγS complex showing representative side (left) and top (right) views (scale bar = 100Å). **(D)** Overlay of low-pass filtered hexamer map and high-resolution sharpened map colored by domain as in (A). **(E)** Final 2.9 Å-resolution sharpened map and **(F)** molecular model colored by individual protomers (P1-P6), with substrate polypeptide (yellow) positioned in the channel. See also Figure S1, S2, and Movie S1.

The ANK-AAA+ domain combination is a unique feature of Skd3. Both the ANK and NBD are required for Skd3 ATPase and disaggregase activity as deletion of either domain ablates activity (Cupo and Shorter, 2020b). How the ANK and NBD collaborate to power disaggregation is unknown. The ANK is comprised of two ankyrin repeats, a linker region, and two more ankyrin repeats (Figure 1A). Ankyrin repeats exhibit a helix-turn-helix conformation, are widely found in nature, and can be adapted for specific protein-protein interactions (Kohl et al., 2003; Mosavi et al., 2004; Parra et al., 2015). Intriguingly, ankyrin repeats are a core component of an ATP-independent disaggregase, cpSRP43 (Jaru-Ampornpan et al., 2013; Jaru-Ampornpan et al., 2010). The NBD of Skd3 is homologous to NBD2 of Hsp104 and ClpB (Cupo and Shorter, 2020b; Erives and Fassler, 2015). Like Hsp104, Skd3 couples ATP hydrolysis to protein disaggregation, which requires conserved AAA+ motifs such as Walker A, Walker B, and pore-loop tyrosines (Cupo and Shorter, 2020b). However, the Skd3 NBD contains an apomorphic insertion at residues L507-I534 that is not observed in any other AAA+ protein (Cupo and Shorter, 2020b; Erzberger and Berger, 2006). What role this insertion plays in Skd3 activity is unknown. Skd3 has an extended CTD that is patterned with both acidic and basic residues (Cupo and Shorter, 2020b). By contrast, *S. cerevisiae* Hsp104 has an extended, acidic CTD that contributes to hexamerization (Mackay et al., 2008). The contribution of the CTD to Skd3 function is also unknown.

SCN-linked mutations in Skd3 cluster in the NBD (Warren et al., 2022), whereas biallelic MGCA-7-linked mutations are scattered throughout all Skd3 domains (Wortmann et al., 2015). SCN-linked mutations impair ATPase and disaggregase activity (Warren et al., 2022), whereas MGCA7-linked mutations impair disaggregase activity in a manner that predicts disease severity (Cupo and Shorter, 2020b). However, MGCA7-linked mutations do not always impair ATPase activity (Cupo and Shorter, 2020b). It is not understood why SCN-linked mutations are dominant-negative, whereas MGCA7-linked mutations are recessive.

It is often assumed that the Skd3 structure and mechanism closely resemble that of bacterial ClpB and yeast Hsp104 (Capo-Chichi et al., 2015; Kanabus et al., 2015; Saunders et al., 2015). Yet, there have been few studies of Skd3 disaggregase activity (Cupo and Shorter, 2020b; Mroz et al., 2020; Warren et al., 2022; Wortmann et al., 2021). Unlike Hsp104 and ClpB, Skd3 does not require Hsp70 or Hsp40 to disaggregate disordered aggregates (Cupo and Shorter, 2020b). Moreover, Skd3 shares only ∼20% sequence identity with *S. cerevisiae* Hsp104 and *E. coli* ClpB and has only one domain, the NBD, in common with Hsp104 and ClpB (Cupo and Shorter, 2020b; Erives and Fassler, 2015). Notably, despite sharing ∼43% identity, even Hsp104 and bacterial ClpB are mechanistically distinct with respect to disaggregase activity (DeSantis et al., 2012; DeSantis et al., 2014; Sweeny and Shorter, 2016). Here, we probe Skd3 structure and function using cryo-electron microscopy (cryo-EM) and mechanistic biochemistry. We report the first high-resolution structure of Skd3 bound to substrate. We also uncover unique mechanistic characteristics that differentiate Skd3 from other disaggregases. Using a mutant subunit doping strategy, we reveal how Skd3 subunits collaborate to drive protein disaggregation. We establish that SCN-linked mutant subunits sharply inhibit Skd3 disaggregase activity, whereas MGCA7-linked mutant subunits do not. Our studies clarify Skd3 structure and mechanism, explain SCN and MGCA7 inheritance patterns, and suggest therapeutic strategies.

## Results

### Structure of _PARL_Skd3 reveals a substrate-bound AAA+ spiral and flexible ANKs

To capture a substrate-bound state of Skd3, we included the model substrate casein, which binds to wild-type (WT) PARL-protease-activated Skd3 (_PARL_Skd3; Figure 1A) (Cupo and Shorter, 2020b). _PARL_Skd3 binding to FITC-labeled casein was determined under different nucleotide conditions (Figure S1A). While binding was identified under all conditions, _PARL_Skd3 bound FITC-casein more effectively in the presence of non-hydrolyzable AMP-PNP (*K_d_*∼0.1µM), ATP (*K_d_*∼0.5µM), or slowly hydrolyzable ATPγS (*K_d_*∼0.4µM) in contrast to ADP or the absence of nucleotide (Figure S1A). Thus, _PARL_Skd3 differs from Hsp104, where only ATPγS facilitates avid polypeptide binding (Gates et al., 2017; Weaver et al., 2017).

Next, we assessed the oligomeric state of _PARL_Skd3 by size-exclusion chromatography (SEC) (Figure 1B, S1B-D). Following incubation with ATPγS, AMP-PNP, or ADP without FITC-casein substrate, _PARL_Skd3 exhibits a broad elution profile with peaks that likely correspond to dodecameric (792 kDa) and hexameric (396 kDa) species, as well as smaller oligomeric or monomeric species (66kDa; Figure S1B). By contrast, in the presence of FITC-casein, _PARL_Skd3 elution shifted toward larger, dodecameric species in all nucleotide conditions (Figure 1B, S1C). Thus, substrate binding by _PARL_Skd3 promotes oligomerization to species larger than the expected hexameric form, likely stabilizing a dodecamer.

Previous structures of ClpB and Hsp104 utilized ATPγS to stabilize substrate-bound states (Gates et al., 2017; Rizo et al., 2019). Indeed, we found that SEC-purified _PARL_Skd3:casein form stable complexes in the presence of ATPγS (Figure 1C, S1E). Reference-free 2D class averages show a variety of top and side views with well-resolved features (Figure 1C, S1E). Top views revealed two classes of particles: a major class with a hexameric-ring structure containing density in the central channel, and a minor class with a heptameric ring and an empty channel (Figure S1E). Top views of the hexameric ring appeared similar to substrate-bound Hsp104 or ClpB (Gates et al., 2017; Rizo et al., 2019), whereas side views exhibited a distinct arrangement with two to three bands of density, indicating a stacked-ring arrangement of the _PARL_Skd3:casein complex (Figure 1C, S1E). Notably, side views show primarily one strong band of density with well-resolved features, whereas the other bands are more diffuse, indicating flexibility or differential occupancy (Figure 1C).

Following 3D classification with four classes, we identified three distinct oligomeric arrangements: a hexameric double-ring complex that contains density in the channel (Class 1), a hexameric three-ring complex that contains density in the central channel for one well-resolved ring (Class 2), and a heptameric form containing two rings and an empty central channel (Class 3) (Figure S1F). Given the low abundance of the heptameric ring and the absence of density for substrate, Class 3 was not pursued further. Class 1 contained the highest percentage of particles (41%) and a well-defined AAA+ ring. Therefore, refinement was performed with this class, resulting in a final overall resolution of 2.9 Å for the _PARL_Skd3:casein complex (Figure S2A-H, Table S1, Movie S1). A molecular model for the hexameric ring comprised of the AAA+ NBD, which refined to the highest resolution (∼2.5 Å) in the map (Figure S2E), was determined using homology models generated by SWISS-Model (Waterhouse et al., 2018).

At an increased threshold, lower-resolution density extends from the N-terminal face of the NBDs and forms a second ring of globular structures that appear separated and flexible (Figure 1D). These separated regions contrast with the extensive contact interfaces made by the NBDs that form the AAA+ ring (Figure 1D). Based on the molecular model of the NBDs and the position of the N-terminal AAA+ residues, we conclude that these separated regions are the N-terminal ANKs (Figure 1A, D). Given this architecture for a hexameric arrangement, we propose that the three-ring structures identified in the 2D class averages and in Class 2 in the 3D classification are likely dodecamers comprised of two Skd3 hexamers that interact via the ANKs, which together form the middle ring of density (Figure 1C; Figure S1F). Considering the flexibility of the ANKs and the second AAA+ ring, it is unclear whether Class 1 is exclusively a hexamer or whether it contains dodecamer particles that are more flexible and not visible in the reconstruction. Indeed, when 2D classification of the particles in Class 1 is performed, weak density for a second AAA+ ring is identified in certain class averages (Figure S2A). Together with the SEC data identifying that _PARL_Skd3 forms a larger, dodecameric species in the presence of substrate and nucleotide (Figure 1B, S1C, D), these data suggest that the active, substrate-bound form of _PARL_Skd3 likely exists in a dynamic equilibrium between hexamer and dodecamer forms.

The NBDs of _PARL_Skd3 adopt a right-handed spiral, wherein 5 protomers directly contact the substrate polypeptide along a 40 Å-length of the channel (Figure 1E, F, Movie S1). These well-resolved protomers (P1-P5) are positioned in a helical arrangement, each with a rise of ∼6 Å and rotation of ∼60° along the substrate. Protomer P6 is at the seam interface between the lowest (P1) and highest (P5) substrate contact sites but is disconnected and has lower resolution, resulting in an asymmetric position within the spiral (Figure 1E, F). This architecture is similar to other substrate-bound AAA+ structures, including Hsp104 and bacterial ClpB (Gates et al., 2017; Rizo et al., 2019). Additional density is also identified in protomers P3-P5 that extends from the small sub-domain of the NBD toward the adjacent clockwise protomer, and likely corresponds to the CTD (Figure 1D).

### Substrate contacts and NBD occupancy support a conserved, stepwise translocation model

Pore loop-substrate interactions and nucleotide states were characterized in the _PARL_Skd3 hexamer structure to elucidate the translocation mechanism. An extended polypeptide is well resolved in the _PARL_Skd3 channel and modeled as a 14-residue poly-A peptide (Figure 2A). Based on our previous structural studies (Gates et al., 2017; Lopez et al., 2020; Rizo et al., 2019) and binding data (Figure S1A, D), we conclude that this extended polypeptide is a nonspecific portion of the incubated FITC-casein. The canonical pore loops (residues 429-432) for protomers P1-P5 extend and directly bind the substrate backbone via the conserved YV motif (Y430 and V431; Figure 2A, B). These pore loops form a spiral staircase of contacts and comprise the primary substrate-binding sites identified in the structure, supporting their established requirement for translocase function (Shorter and Southworth, 2019). Indeed, mutation of the conserved tyrosine to alanine (Y430A) reduces ATPase activity and abolishes disaggregase activity (Cupo and Shorter, 2020b). We now find that V431G also reduces _PARL_Skd3 ATPase activity and abolishes disaggregase activity (Figure 2C,D). These results are consistent with our structure of _PARL_Skd3, which identifies direct substrate contact by Y430 and V431.

**Figure 2.**
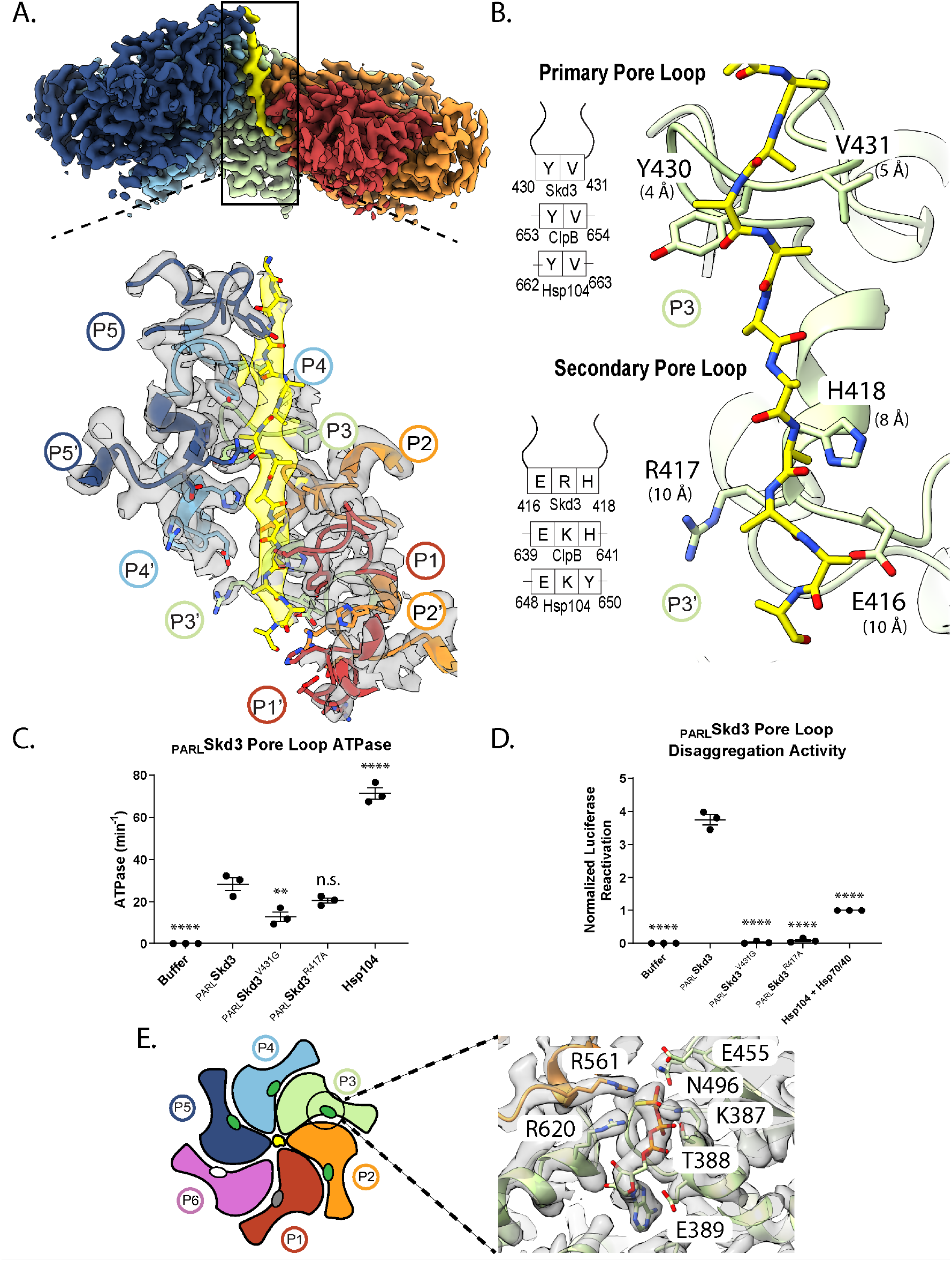
Spiral of pore loop-substrate contacts and nucleotide states of _PARL_Skd3. **(A)** Cryo-EM density map (top) of protomers (P1-P5) and substrate (yellow) and expanded map plus model view (bottom) of the channel including the primary (P1-P5) and secondary (P1’-P5’) loops interacting in a spiral along the 14-residue substrate strand. P1 is the canonical pore loop from protomer 1 and P1’ is the secondary pore loop from protomer 1. **(B)** The primary and secondary pore loop-substrate contacts for P4, including distances to substrate (measured between α-carbons) and a schematic indicating conservation among disaggregases. **(C)** ATPase activity of _PARL_Skd3, _PARL_Skd3^V431G^, _PARL_Skd3^R417A^, and Hsp104. ATPase activity was compared to _PARL_Skd3 using one-way ANOVA and a Dunnett’s multiple comparisons test (N = 3, individual data points shown as dots, bars show mean ± SEM, **p<0.01, ****p<0.0001). **(D)** Luciferase disaggregase activity of _PARL_Skd3, _PARL_Skd3^V431G^, _PARL_Skd3^R417A^, and Hsp104 plus Hsp70 and Hsp40. Luciferase activity was buffer subtracted and normalized to Hsp104 plus Hsp70 and Hsp40. Disaggregase activity was compared to _PARL_Skd3 using one-way ANOVA and a Dunnett’s multiple comparisons test (N = 3, individual data points shown as dots, bars show mean ± SEM, ****p<0.0001). **(E)** Schematic indicating nucleotide states (ovals) for each protomer (ATP = Green; ADP = grey; apo = white) and expanded view of the map plus model P4 nucleotide pocket showing density for ATP and conserved interacting residues (including Arg finger (R561), sensor-1 (N496), sensor-2 (R620), Walker A (K387) and Walker B (E455)) that define the ATP state. See also Figure S2 and Movie S1.

An additional spiral of substrate interactions at the channel exit is formed by secondary pore-loop motifs from protomers P2-P5 (Figure 2A, B). In _PARL_Skd3, residues E416, R417 and H418 comprise this secondary pore loop, which is positioned in line with canonical YV loops above, but slightly further away (∼9 Å) from the substrate backbone (Figure 2A, B). To further characterize the role of the secondary pore loop, we generated _PARL_Skd3^R417A^, which exhibited similar ATPase activity to _PARL_Skd3 (Figure 2C), but diminished disaggregase activity (Figure 2D). This loss of function is substantially more severe than that caused by equivalent mutations in ClpB or Hsp104 (Howard et al., 2020; Rizo et al., 2019). Thus, the secondary pore loops play a more critical role in _PARL_Skd3 disaggregase activity than in Hsp104 or ClpB.

The nucleotide-binding pockets in the substrate-bound _PARL_Skd3 complex are positioned at the inter-protomer interfaces with conserved AAA+ residues contacting nucleotide (Figure 2E, S2I). For protomers P2-P5, these pockets are well resolved, revealing a bound ATP molecule that is contacted by canonical Walker A (K387), Walker B (E455), sensor-1 (N496), and sensor-2 (R620) residues (Figure 2E, S2I). The Arg-finger residue (R561) is provided by the neighboring clockwise protomer, and positioned one step lower along the substrate, contacting the γ-phosphate of ATP in protomers P3-P5 (Figure 2E, S2I). For protomer P2, complete density for ATP is present, but the Arg-finger residue from P1 is positioned further away and not in contact, indicating a potential intermediate state (Figure S2I). Notably, these ATP-bound states are only found for protomers that contact substrate (Figure 2A, S2I). Conversely, density for nucleotide is more poorly resolved in protomers P1 and P6 at the spiral seam (Figure 2A, S2I). Nucleotide appears absent from P6, indicating an apo state, whereas P1 is likely ADP-bound. Thus, post-hydrolysis states likely coincide with substrate release at the seam. These findings indicate that _PARL_Skd3 employs a conserved hydrolysis cycle similar to other AAA+ disaggregases and translocases (Gates et al., 2017; Puchades et al., 2017; Rizo et al., 2019). Based on this model, ATP hydrolysis and substrate release occur at the lower contact sites in the spiral (P1), whereas ATP binding promotes substrate re-binding to the top position (P5) along the substrate, enabling a rotary mechanism involving two amino steps along the substrate during processive translocation (Shorter and Southworth, 2019). However, other kinetic paths or non-processive events may also be possible (Durie et al., 2019; Fei et al., 2020).

### ANKs mediate _PARL_Skd3 dodecamer formation and enable disaggregase activity

The Skd3 ANK is a unique feature among AAA+ unfoldases and is required for Skd3 disaggregase activity, indicating a distinct functional role (Cupo and Shorter, 2020b). Cryo-EM of _PARL_Skd3^NBD^, which lacks the ANK domain, reveals well-resolved single hexamers but no larger oligomers (Figure S3A). Based on our structural analysis, the ANK forms a middle ring of interactions that support a double hexamer (dodecamer) arrangement of the complex (Figure 1C). Thus, the ANK is not required for hexamerization, but is important for stabilizing the larger dodecamer state. SEC indicates that this dodecameric form is likely the predominant species in the presence of substrate (Figure 1B, S1C). However, the dodecamer is less well-represented following 2D and 3D cryo-EM analysis, with ∼15% of particles possessing the three-ring architecture of Class 2 (Figure S1F). Moreover, flexibility of the ANKs and the tilted arrangement of the AAA+ rings likely limit structure determination of the full dodecamer complex from Class 2. Nonetheless, two full Skd3 hexamer models could be docked into the low-resolution Class 2 map, revealing that the ANKs mediate contacts across the two hexamers (Figure S3B, Table S1). Resolution of the second NBD hexamer was insufficient to identify substrate in the channel or the spiral protomer arrangement. Conversely, in addition to the high-resolution AAA+ ring, the final map of the hexamer class (Class 1) contains strong globular density extending from the N-terminal face that is consistent with the helical bundles of ankyrin repeats (Figure 1D). Therefore, analysis of the complete hexamer arrangement was further pursued with the Class 1 map.

Structural information for the Skd3 ANK is not available. Thus, we used the Alpha-fold structure prediction to determine a model for the ANK (Jumper et al., 2021). This secondary structural model is predicted with high confidence based on the pLDDT score and low predicted align error values (Figure S3C, D). The confidence was highest in both the ANK and NBD domains (Figure S3D). Based on the Alpha-fold model the ANK is predicted to adopt four two-helix bundle structures that match canonical ANKs (Figure 3A). Starting at the N-terminus, this structure consists of two ankyrin repeats (1 and 2), a 66-residue linker (L) that is mostly disordered, and two additional ankyrin repeats (3 and 4) (Figure 3A). Curiously, the linker is the exact length of two ankyrin repeats and appears to have some cryptic elements of an ankyrin repeat within its primary sequence (Figure S3F). Alpha-fold predicts some helical regions within the linker, and these regions partially align to the other repeats (Figure 3A, S3C,D). Thus, the linker region may impart some ankyrin-like functions to Skd3. Notably, repeat 4 forms an extended helix that transitions directly into the N-terminal region of the NBD without a separate linker between the domains (Figure 3A). This continuous helix likely adds some stability to the position of the ANKs given that inter-protomer contacts are not present in the ANK ring. The four ankyrin repeats bundle together in the Alpha-fold model and dock well into the globular density adjacent to the NBD (Figure 3B, Movie S1). The density for the ANK is more prominent for protomers P2-P5, which are bound to substrate and better resolved compared to the spiral seam (Figure 3B). To further resolve the ANK, focus classification was performed on the P3 ANK. Resulting classes reveal the ANK adopts different positions, indicating the flexibility of the ankyrin-repeat 4/NBD connecting helix (Figure 3C, D, S3E). Notably, Class 1 contains additional density that projects from the globular ANKs towards the central channel and may correspond to the linker based on our molecular model (Figure S3E).

**Figure 3.**
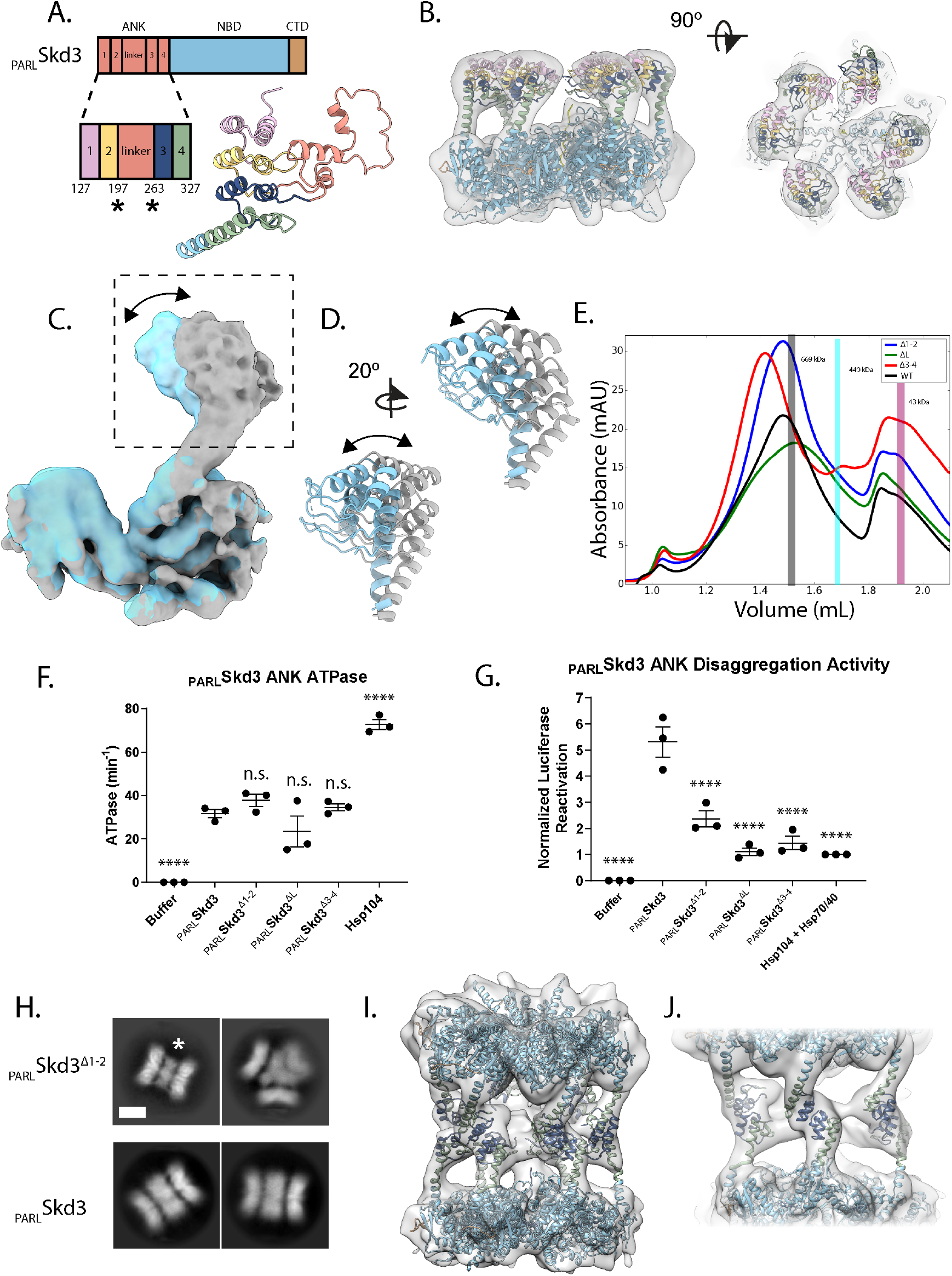
Ankyrin repeats mediate head-to-head contacts required for dodecamer formation. **(A)** Schematic and predicted model of _PARL_Skd3 ANK, colored based repeat number and linker. **(B)** Side (left) and top (right) views of the filtered Class 1 map and docked model, colored as in (A), identifying ANK position. **(C, D)** Overlay of two different classes resolved from focus classification around P3 identifying rotation of the globular ANK region (arrow). **(E)** SEC of _PARL_Skd3 (black), _PARL_Skd3^Δ1-2^ (blue), _PARL_Skd3^ΔL^ (green), and _PARL_Skd3^Δ3-4^ (red) incubated with ATPγS and casein. Vertical bars indicate elution position of molecular-weight standards: thyroglobulin (669 kDa), ferritin (440 kDa), and ovalbumin (43 kDa), and are representative of _PARL_Skd3 dodecamer (grey), hexamer (cyan), or monomer (magenta) size. **(F)** ATPase activity of _PARL_Skd3, _PARL_Skd3^Δ1-2^, _PARL_Skd3^ΔL^, _PARL_Skd3^Δ3-4^, and Hsp104. ATPase activity was compared to _PARL_Skd3 using one-way ANOVA and a Dunnett’s multiple comparisons test (N = 3, individual data points shown as dots, bars show mean ± SEM, ****p<0.0001). **(G)** Luciferase disaggregase activity of _PARL_Skd3, _PARL_Skd3^Δ1-2^, _PARL_Skd3^ΔL^, _PARL_Skd3^Δ3-4^, and Hsp104 plus Hsp70 and Hsp40. Luciferase activity was buffer subtracted and normalized to Hsp104 plus Hsp70 and Hsp40. Disaggregase activity was compared to _PARL_Skd3 using one-way ANOVA and a Dunnett’s multiple comparisons test (N = 3, individual data points shown as dots, bars show mean ± SEM, ****p<0.0001). **(H)** 2D class averages comparing _PARL_Skd3^Δ1-2^ (top) and _PARL_Skd3 (bottom) oligomers with middle band of ANK density indicated (*). Note the triple-hexamer arrangement (top right) is only identified for _PARL_Skd3^Δ1-2^. Scale bar, 100Å. **(I, J)** Dodecamer map and model of _PARL_Skd3_Δ1-2_ colored by individual domains showing model for ANK 3,4 interactions across hexamers. See also Figure S3 and Movie S2.

To assess the contribution of specific regions of the ANK toward Skd3 functionality, we generated _PARL_Skd3 variants with ankyrin repeat 1 and 2 deleted (ΔY127-G196, _PARL_Skd3^Δ1-2^), the linker deleted (ΔD197-A262, _PARL_Skd3^ΔL^), or ankyrin repeats 3 and 4 deleted (ΔS263-K325,_PARL_Skd3^Δ3-4^) (Figure S3G). In the presence of casein and ATPγS, _PARL_Skd3^Δ1-2^ and _PARL_Skd3^Δ3-4^ formed predominantly dodecamers rather than hexamers like _PARL_Skd3 (Figure 1B, 3E). By contrast, _PARL_Skd3^ΔL^ exhibited reduced dodecamer formation, and was shifted more toward the hexameric form (Figure 3E). Unlike _PARL_Skd3^NBD^, which exhibits reduced ATPase activity (Cupo and Shorter, 2020b), _PARL_Skd3^Δ1-2^, _PARL_Skd3^ΔL^, and _PARL_Skd3^Δ3-4^ exhibited similar ATPase activity to _PARL_Skd3 (Figure 3F). Thus, a portion of the N-terminal ANK is required to maintain _PARL_Skd3 ATPase activity. By contrast, _PARL_Skd3^Δ1-2^, _PARL_Skd3^ΔL^, and _PARL_Skd3^Δ3-4^ exhibited reduced disaggregase activity (Figure 3G), indicating that the ANK enables _PARL_Skd3 to couple ATP hydrolysis to protein disaggregation. Deletion of the linker had the largest effect (Figure 3G). Importantly, _PARL_Skd3^ΔL^ is impaired in dodecamer formation in the presence of substrate (Figure 3E). Thus, dodecamer formation may promote disaggregase activity. These findings suggest that the ANKs may play multiple roles in protein disaggregation, including dodecamerization, potentially supported by the linker, and possible direct roles in substrate binding mediated by ankyrin repeats 1-4. Initial EM analysis revealed that _PARL_Skd3^Δ1-2^ forms more stable dodecamers compared to _PARL_Skd3^ΔL^ and _PARL_Skd3^Δ3-4^. Thus, _PARL_Skd3^Δ1-2^ was investigated further by cryo-EM. 2D averages of _PARL_Skd3^Δ1-2^ show a well-resolved middle ring of ANKs that is smaller in diameter than _PARL_Skd3 (Figure 3H). 3D classification of _PARL_Skd3^Δ1-2^ identified two distinct oligomeric forms. Class 1 and Class 2 are dodecamers with different relative positions of the AAA+ rings, whereas Class 3 is a trimer of hexamers (Figure S3H, I). Refinement of Class 1 was pursued due to the more homogeneous arrangement of the central ANK ring and improved density for the second AAA+ ring compared to the _PARL_Skd3 complex (Figure 3I, J, S3J-M). Whereas the overall resolution was low (∼9Å), a dodecameric model with ankyrin repeats 3 and 4 fit well into the density, and revealed head-to-head ANK contacts around the central ring (Figure 3I, J, Table S1, Movie S2). These results further support that the ANK interacts in a head-to-head manner to mediate dodecamer formation. Given that _PARL_Skd3^Δ3-4^ can also form the dodecamer as can _PARL_Skd3^ΔL^ to a lesser extent (Figure 3E), these findings indicate plasticity in how the ANK mediates cross-contacts to form the dodecamer. Based on these results we suggest that deleting specific ankyrin repeats or the linker reduces this interactive plasticity and thereby reduces disaggregase activity (Figure 3G).

### A unique insertion within the _PARL_Skd3 NBD regulates the AAA+ motor

Skd3 contains an insertion (residues L507-I534) within the NBD that is highly conserved across Skd3 homologues but is not observed in Hsp104 or other HCLR class AAA+ proteins (Figure 4A, S4A) (Cupo and Shorter, 2020b). Based upon our _PARL_Skd3 reconstruction, we modeled part of the insertion, but 17 residues (517-533) were unaccounted for (Figure 4B, S4B). When compared to Hsp104, the insertion in Skd3 extends past the loop that is present in Hsp104 (Figure S4C) and protrudes from the hexamer exterior (Figure 4B). Purified _PARL_Skd3^ΔL507-I534^ formed a large oligomeric species that is not observed for _PARL_Skd3 (Figure S4D). However, upon addition of casein, _PARL_Skd3^ΔL507-I534^ shifts to predominantly dodecamers (Figure S4D). Indeed, in the presence of casein, _PARL_Skd3^ΔL507-I534^ shifted more toward dodecamers than hexamers compared to _PARL_Skd3 (Figure 1B, S1C, D, S4D). _PARL_Skd3^ΔL507-I534^ exhibited elevated ATPase and disaggregase activity compared to _PARL_Skd3 (Figure 4C, D). These findings suggest that the L507-I534 insertion acts as a regulatory element, which slows _PARL_Skd3 ATPase activity and tunes disaggregase activity. The location of the L507-I534 insertion on the exterior of the hexamer could enable it to serve as a site for regulatory factors to bind or post-translationally modify Skd3.

**Figure 4.**
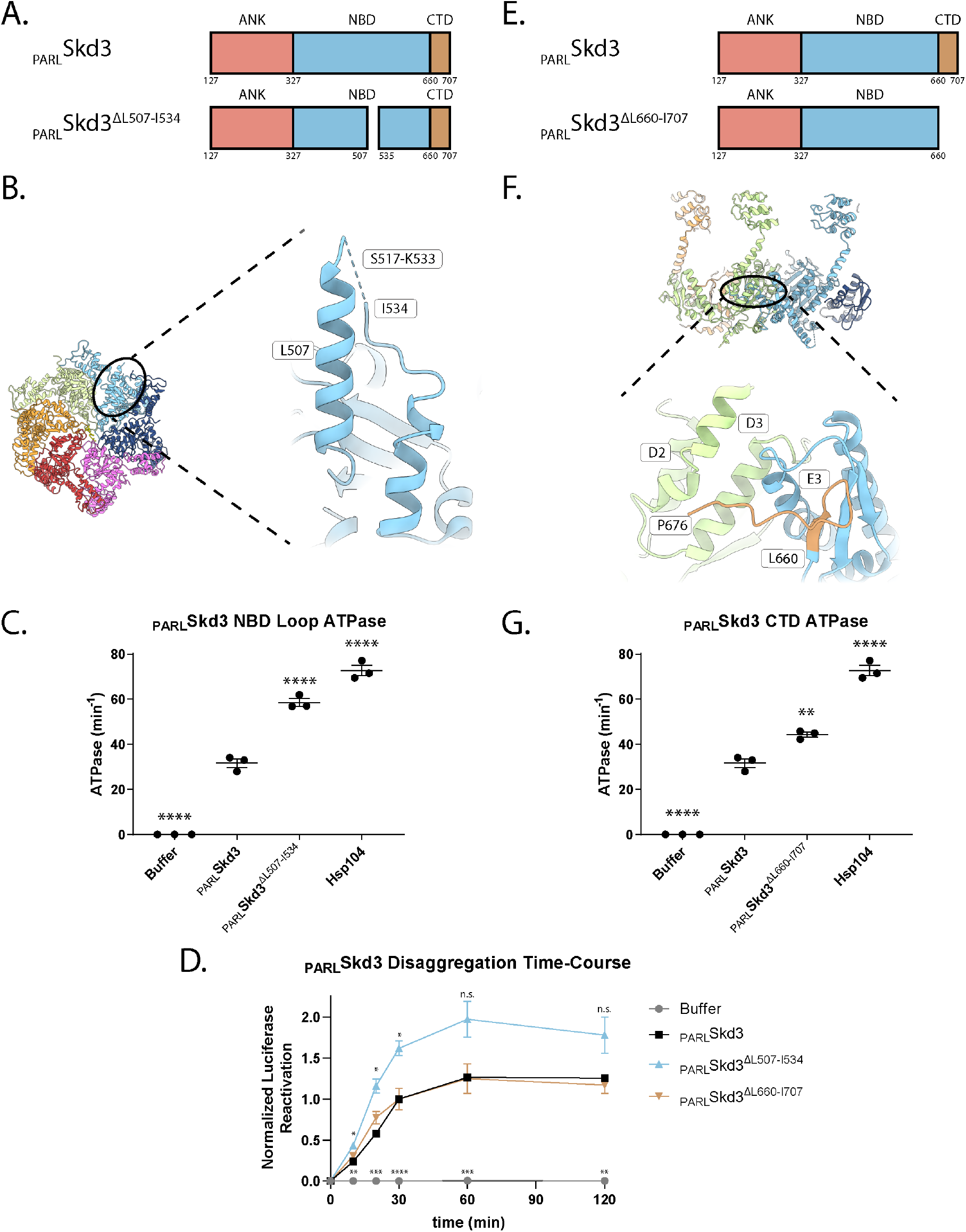
An NBD insertion and the CTD regulate _PARL_Skd3 activity. **(A)** Domain schematic of _PARL_Skd3 and _PARL_Skd3^ΔL507-I534^ colored by individual domains. **(B)** Top view of _PARL_Skd3 hexamer structure and expanded view of NBD-insertion region residues 449-552 (residues 517-533 are not resolved (dashed line)) in P4. **(C)** ATPase activity of _PARL_Skd3, _PARL_Skd3^ΔL507-I534^, and Hsp104. Data are from the same experiments as Figure 3F. ATPase activity was compared to _PARL_Skd3 using one-way ANOVA and a Dunnett’s multiple comparisons test (N = 3, individual data points shown as dots, bars show mean ± SEM, ****p<0.0001). **(D)** Luciferase disaggregation time course showing that _PARL_Skd3^ΔL507-I534^ has accelerated disaggregase activity at some time points whereas _PARL_Skd3^ΔL660-I707^ does not. Luciferase activity was buffer subtracted and normalized to _PARL_Skd3 30 min time point. Luciferase activity was compared to _PARL_Skd3 using one-way ANOVA and a Dunnett’s multiple comparisons test (N = 3, individual data points shown as dots, bars show mean ± SEM, **p<0.01, ***p<0.001, ****p<0.0001). **(E)** Domain schematic of _PARL_Skd3 and _PARL_Skd3^ΔL660-I707^ colored by individual domains. **(F)** Side view of _PARL_Skd3 hexamer structure and expanded view of P3-P4 with the P4 CTD model (brown) shown adjacent P3 with potential interacting helices indicated. **(G)** ATPase activity of _PARL_Skd3 and _PARL_Skd3^ΔL660-I707^. Data are from the same experiments as Figure 3F. ATPase activity was compared to _PARL_Skd3 using one-way ANOVA and a Dunnett’s multiple comparisons test (N = 3, individual data points shown as dots, bars show mean ± SEM, **p<0.01, ****p<0.0001). See also Figure S4.

### Deletion of the _PARL_Skd3 CTD mildly stimulates ATPase activity

Deletion of the extended, acidic CTD of *S. cerevisiae* Hsp104 results in hexamerization defects (Mackay et al., 2008). Like Hsp104 and in contrast to other HCLR clade AAA+ proteins, Skd3 has an extended CTD from residues 660 to 707 (Figure 4E) (Cupo and Shorter, 2020b). Unlike Hsp104, however, the Skd3 CTD is patterned with acidic and basic residues, whereas the Hsp104 CTD is acidic (Cupo and Shorter, 2020b). From the reconstruction of _PARL_Skd3, 14 residues of the CTD were evident in protomers P3-P5 (Figure 1D). These residues fit along the side of the adjacent protomer and are near helices D2 and D3 (Figure 4F, S2F). Residues within ∼4 Å of the CTD include E340 and Q341 in D2, and R362 in D3. Additional contacts could occur with helix E3 of the same protomer (Figure 4F, S2F). We purified _PARL_Skd3 lacking the CTD (_PARL_Skd3^ΔL660-I707^), which formed hexamers and dodecamers similar to _PARL_Skd3 (Figure 1A, S4G). However, _PARL_Skd3^ΔL660-I707^ exhibited mildly increased ATPase activity (Figure 4H), whereas disaggregase activity was similar to _PARL_Skd3 (Figure 4D). Thus, the CTD enables efficient coupling of _PARL_Skd3 ATPase activity to disaggregase activity. Overall, these findings suggest that the _PARL_Skd3 CTD plays a different role than the Hsp104 CTD.

### _PARL_Skd3 is functional at low ATP concentrations

Mitochondria maintain lower ratios of ATP:ADP and overall lower ATP concentrations than the cytoplasm (Gellerich et al., 2002; Heldt et al., 1972; Imamura et al., 2009). To determine how _PARL_Skd3 might operate under a variety of nucleotide conditions, we next established that _PARL_Skd3 ATPase activity has a V_max_ of ∼24min^-1^ and a K_M_ of ∼65μM (Figure 5A). This K_M_ is similar to the value reported for _MPP_Skd3 (Mroz et al., 2020). Thus, removal of the inhibitory peptide by PARL does not grossly alter K_M_. By contrast, the K_M_ of Hsp104 is ∼5-11mM (Grimminger et al., 2004; Schirmer et al., 1998). Strikingly, _PARL_Skd3 can also maintain maximal disaggregase activity at low ATP concentrations (Figure 5B). _PARL_Skd3 maintained ∼50% disaggregase activity at the lowest concentration of ATP tested (0.434 mM) (Figure 5B). Thus, _PARL_Skd3 is likely adapted to operate effectively at lower ATP concentrations than Hsp104. Hsp104 is sharply inhibited by mixing ADP with ATP (Grimminger et al., 2004; Hattendorf and Lindquist, 2002; Klosowska et al., 2016). Indeed, even a 5:1 ATP:ADP ratio can diminish Hsp104 activity (Klosowska et al., 2016). To investigate the effect of ADP on _PARL_Skd3, we assessed _PARL_Skd3 disaggregase activity under different ATP:ADP ratios while keeping the total nucleotide concentration constant. Under these conditions, ADP did not affect luciferase activity, indicating that any effects of ADP reflect direct effects on _PARL_Skd3. _PARL_Skd3 is inhibited by ADP, but maintains ∼50% activity at a 5:1 ATP:ADP ratio (Figure 5C), which can inactivate Hsp104 (Klosowska et al., 2016). The half-maximal inhibitory concentration (IC_50_) of ADP at a constant concentration of ATP (5mM) was ∼1.2mM (Figure 5D). These findings suggest that _PARL_Skd3 is less sensitive than Hsp104 to inhibition by ADP. Indeed, _PARL_Skd3 is likely adapted to function at the lower ATP:ADP ratios found in mitochondria.

**Figure 5.**
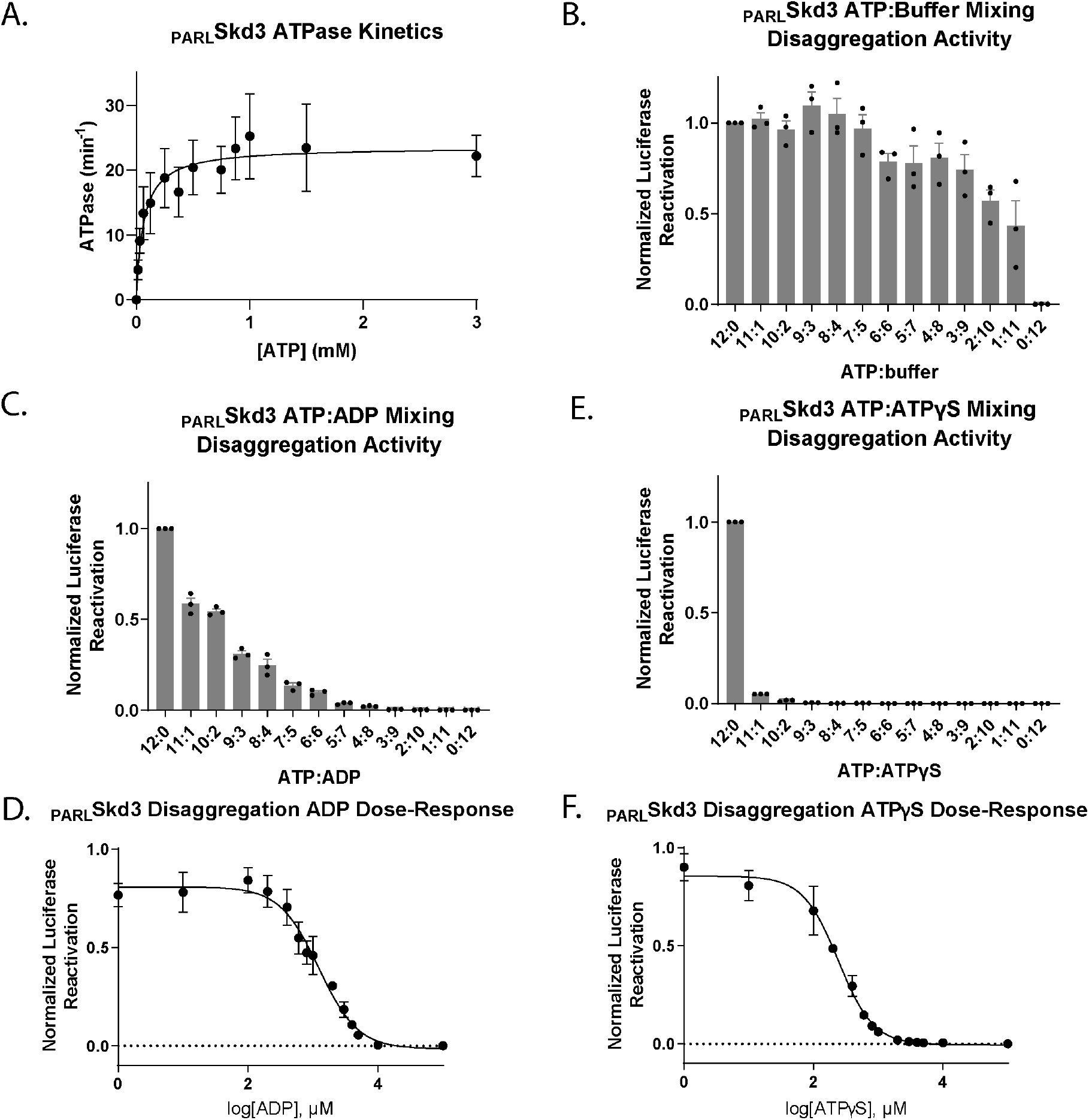
_PARL_Skd3 is functional at low ATP concentrations. **(A)** Michaelis-Menten plot of _PARL_Skd3 ATPase activity. V_max_ was determined to be ∼23.6 min^-1^. K_M_ was determined to be ∼64.6μM (N = 3, data shown as mean ± SEM). **(B)** Luciferase disaggregase activity of _PARL_Skd3 in the presence of various ratios of ATP:buffer. The concentrations used were 12:0 (5mM), 11:1 (4.58mM), 10:2 (4.17mM), 9:3 (3.75mM), 8:4 (3.33mM), 7:5 (2.92mM), 6:6 (2.5mM), 5:7 (2.08mM), 4:8 (1.67mM), 3:9 (1.25mM), 2:10 (0.83mM), 1:11 (0.42mM), and 0:12 (0mM). Disaggregase activity was buffer subtracted and normalized to _PARL_Skd3 plus ATP (N = 3, individual data points shown as dots, bars show mean ± SEM). **(C)** Luciferase disaggregase activity of _PARL_Skd3 in the presence of various ratios of ATP:ADP where total concentration of nucleotide was maintained at 5mM. Disaggregase activity was buffer subtracted and normalized to _PARL_Skd3 plus ATP (N = 3, individual data points shown as dots, bars show mean ± SEM). **(D)** Luciferase disaggregase activity of _PARL_Skd3 in the presence of a constant concentration of ATP (5mM) and increasing concentrations of ADP. Disaggregase activity was buffer subtracted and normalized to _PARL_Skd3 plus ATP (N = 3, data shown as mean ± SEM). **(E)** Luciferase disaggregase activity of _PARL_Skd3 in the presence of various ratios of ATP:ATPγS where total concentration of nucleotide was maintained at 5mM. Disaggregase activity was buffer subtracted and normalized to _PARL_Skd3 plus ATP (N = 3, individual data points shown as dots, bars show mean ± SEM). **(F)** Luciferase disaggregase activity of _PARL_Skd3 in the presence of a constant concentration of ATP (5mM) and increasing concentrations of ATPγS. Disaggregase activity was buffer subtracted and normalized to _PARL_Skd3 plus ATP (N = 3, data shown as mean ± SEM).

### _PARL_Skd3 disaggregase activity is sharply inhibited by ATPγS

Next, we assessed how _PARL_Skd3 disaggregase activity is affected by the slowly hydrolyzable ATP analogue, ATPγS. Like Hsp104, _PARL_Skd3 is inactive in the presence of ATPγS as the sole nucleotide (Cupo and Shorter, 2020b; DeSantis et al., 2012; Doyle et al., 2007; Torrente et al., 2016). However, Hsp104 disaggregase activity against disordered aggregates can be stimulated at specific ratios of ATP:ATPγS (∼3:1-1:5), whereas Hsp104 disaggregase activity against amyloid is invariably inhibited by ATPγS in the presence of ATP (DeSantis et al., 2012). These differences suggest that Hsp104 employs distinct mechanisms of subunit collaboration to disaggregate disordered aggregates versus amyloid (DeSantis et al., 2012). To assess the effect of ATPγS on _PARL_Skd3, we measured _PARL_Skd3 disaggregase activity under different ATP:ATPγS ratios while keeping the total nucleotide concentration constant. Under these conditions, ATPγS did not affect luciferase activity, indicating that any effects of ATPγS reflect direct effects on _PARL_Skd3. _PARL_Skd3 is sharply inhibited by ATPγS (Figure 5E). Even an 11:1 ATP:ATPγS ratio strongly inhibits _PARL_Skd3 (Figure 5E). The IC_50_ of ATPγS at a constant concentration of ATP (5mM) was ∼242µM (Figure 5F). Thus, in contrast to Hsp104 (DeSantis et al., 2012), _PARL_Skd3 disaggregase activity against disordered luciferase aggregates is not stimulated by mixtures of ATP and ATPγS. The distinctive responses of _PARL_Skd3 to ADP and ATPγS reveal key differences in how _PARL_Skd3 and Hsp104 disaggregase activity are regulated. Moreover, our findings suggest that ADP-bound _PARL_Skd3 subunits are less inhibitory than ATPγS-bound _PARL_Skd3 subunits. Indeed, the sharp inhibition of _PARL_Skd3 disaggregase activity by ATPγS indicates that _PARL_Skd3 is sensitive to individual subunits that hydrolyze ATP slowly.

### _PARL_Skd3 is a subglobally cooperative protein disaggregase

Next, to further define mechanochemical coupling mechanisms of _PARL_Skd3, we harnessed a mutant subunit doping strategy to assess the contribution of individual _PARL_Skd3 subunits toward ATPase activity and disaggregase activity. For this purpose, we modeled Skd3 as a hexamer, which forms the functional AAA+ cassette (Figure 2E). In this strategy, mutant _PARL_Skd3 subunits defective in ATP hydrolysis or substrate binding are mixed with WT _PARL_Skd3 subunits to generate heterohexameric ensembles according to a binomial distribution that is determined by the WT:mutant ratio (Figure 6A). As mutant _PARL_Skd3 concentration in the mixture increases, the probability of mutant _PARL_Skd3 incorporation into a _PARL_Skd3 hexamer increases (Figure 6A). This approach has revealed mechanochemical coupling mechanisms of other NTPases, including bacterial ClpB and Hsp104 (DeSantis et al., 2012; DeSantis et al., 2014; Moreau et al., 2007; Shivhare et al., 2019; Sweeny et al., 2015; Torrente et al., 2016).

**Figure 6.**
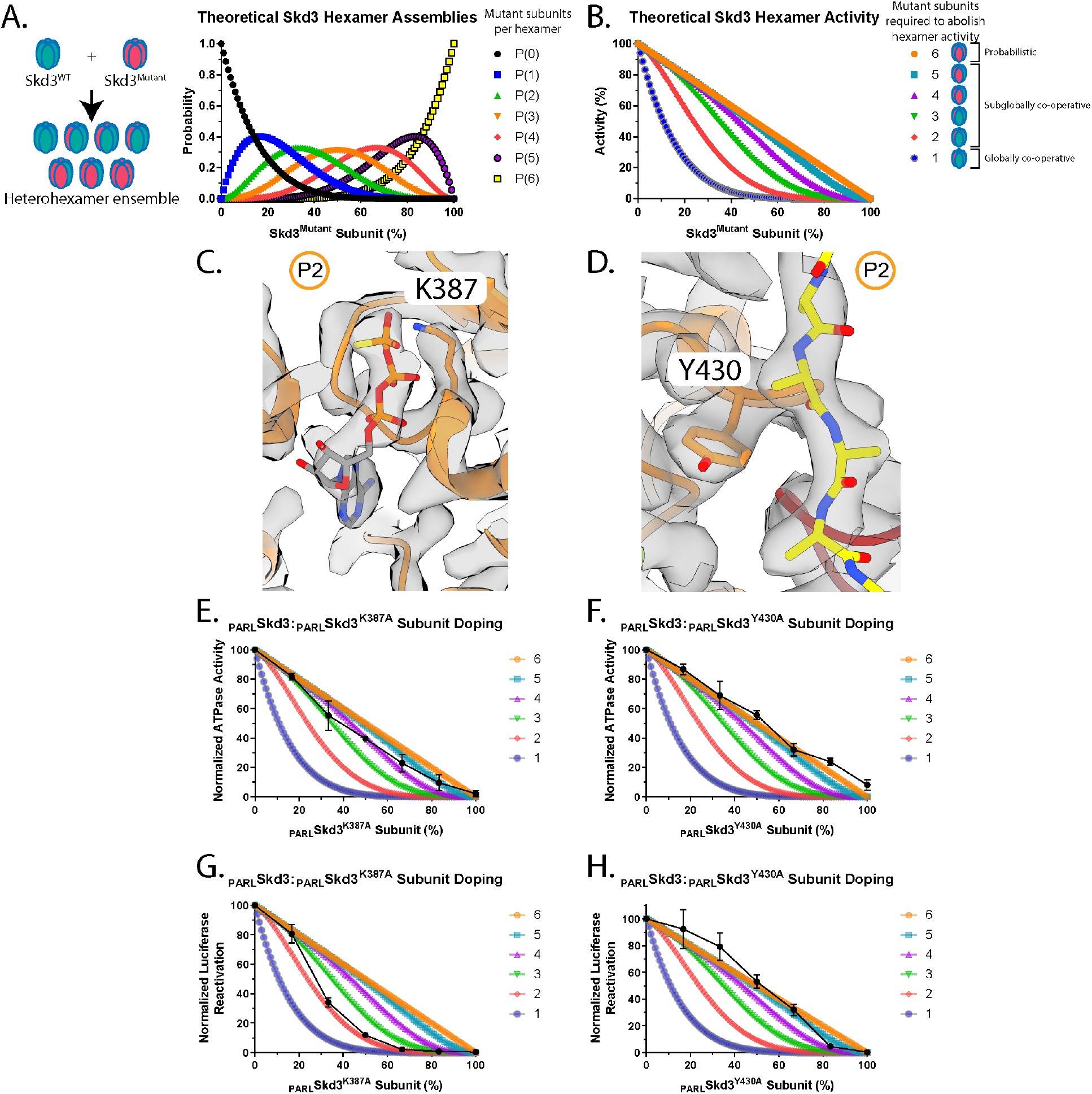
Skd3 is a subglobally cooperative protein disaggregase. **(A)** Theoretical _PARL_Skd3 hexamer ensembles containing zero (black), one (blue), two (green), three (orange), four (red), five (purple), and six (yellow) mutant subunits as a function of the fraction of mutant subunit present. **(B)** Theoretical _PARL_Skd3 activity curves where one or more (blue), two or more (red), three or more (green), four or more (purple), five or more (light blue), or six (orange) mutant subunits are needed to ablate hexamer activity. In a probabilistic model, 6 mutant subunits are required to poison a hexamer. In a subglobally cooperative model, 2-5 mutant subunits are required to poison hexamer activity. In a globally cooperative model, a single mutant subunit is sufficient to poison a hexamer. **(C, D)** Map and model of protomer P2 with nucleotide and residues K387 (C) and Y430 (D) shown, respectively. **(E, F)** ATPase activity of _PARL_Skd3 mixed with various ratios of _PARL_Skd3^K387A^ (E) or _PARL_Skd3^Y430A^ (F). ATPase activity was buffer subtracted and normalized to _PARL_Skd3 (N = 3, data shown as black dots with mean ± SEM). **(G, H)** Luciferase disaggregase activity of _PARL_Skd3 mixed with various ratios of _PARL_Skd3^K387A^ (G) or _PARL_Skd3^Y430A^ (H). Disaggregase activity was buffer subtracted and normalized to _PARL_Skd3 (N = 3, data shown as black dots with mean ± SEM). See also Figure S5.

This strategy depends on robust formation of randomized heterohexamer ensembles, which requires exchange of subunits between WT and mutant _PARL_Skd3 hexamers such that mutant subunits mix equally well into heterohexamers as WT (Figure 6A). To assess subunit mixing, we labeled _PARL_Skd3 with Alexa488 or Alexa594, which can form a Förster resonance energy transfer (FRET) pair. Labeled _PARL_Skd3 retained ATPase and disaggregase activity, indicating that labeling did not eliminate functionality (Figure S5A, B). We mixed Alexa488-labeled _PARL_Skd3 and Alexa594-labeled _PARL_Skd3 in the absence of substrate, where the hexamer is more populated (Figure S1B, C). Thus, FRET likely reflects subunit mixing within the hexamer. For WT _PARL_Skd3, a robust FRET signal was observed within a few minutes, indicating subunit mixing on the minute timescale similar to Hsp104 (Figure S5D) (DeSantis et al., 2012). Importantly, mutant _PARL_Skd3 subunits were effectively incorporated into WT _PARL_Skd3 hexamers (Figure S5D). Thus, _PARL_Skd3^K387A^ (Walker A) subunits likely incorporate into WT _PARL_Skd3 hexamers as effectively as WT _PARL_Skd3 subunits, whereas _PARL_Skd3^Y430A^ (pore loop) subunits incorporated into WT _PARL_Skd3 hexamers ∼16% less effectively than WT _PARL_Skd3 subunits (Figure S5D). These findings indicate that _PARL_Skd3 likely forms dynamic hexamers that exchange subunits on the minute timescale. Moreover, specific mutant _PARL_Skd3 subunits incorporate effectively into WT _PARL_Skd3 hexamers. Thus, _PARL_Skd3 provides a tractable system for mutant doping studies.

This rapid subunit exchange enables formation of _PARL_Skd3 heterohexamer ensembles comprised of WT and mutant subunits according to a binomial distribution that varies as a function of the molar ratio of each subunit (Figures 6A). Using this distribution, we can predict how _PARL_Skd3 activity would be inhibited at various WT:mutant ratios if a specific number of _PARL_Skd3 mutant subunits inactivate the hexamer (Figure 6B). For example, if all six _PARL_Skd3 subunits must work together, then one mutant subunit would abolish hexamer activity (Figure 6B, dark blue curve). At the other extreme, if the activity of a single _PARL_Skd3 subunit within the hexamer is sufficient, then some activity would still be observed with five mutant subunits per hexamer, and only six mutant subunits would abolish activity (Figure 6B, orange line). Thus, by comparing experimental data with theoretical plots, we can determine whether subunit collaboration within _PARL_Skd3 hexamers is probabilistic (6 mutant subunits abolish activity), subglobally cooperative (2-5 mutant subunits abolish activity), or globally cooperative (one mutant subunit abolishes activity).

Next, we titrated _PARL_Skd3 with buffer over the concentration range of the subunit doping ATPase assay and found a linear decline in ATPase activity (Figure S5E). Thus, when titrating mutant _PARL_Skd3, a sharper than linear decline in ATPase activity indicates inhibitory effects of mutant subunits incorporated into hexamers. Similarly, we titrated _PARL_Skd3 with buffer over a range of concentrations to assess disaggregase activity (Figure S5F). We selected saturating _PARL_Skd3 concentrations to ensure that any observed effects on disaggregase activity upon mixing WT and mutant are not caused by a mere decrease in the concentration of WT _PARL_Skd3 (DeSantis et al., 2012; Werbeck et al., 2008).

_PARL_Skd3^K387A^ (Walker A) and _PARL_Skd3^Y430A^ (pore loop) are inactive for ATPase and disaggregase activity (Cupo and Shorter, 2020b). The Walker A residue, K387, contacts the β and γ-phosphate of ATP and mutating this residue to alanine is predicted to reduce ATP binding and hydrolysis (Figure 6C) (Erzberger and Berger, 2006; Hanson and Whiteheart, 2005; Puchades et al., 2020; Wendler et al., 2012). The pore-loop tyrosine, Y430, engages substrate and its mutation to alanine is predicted to reduce substrate binding (Figure 2B, 6D) (Cupo and Shorter, 2020b). We assembled heterohexamer ensembles of _PARL_Skd3 with _PARL_Skd3^K387A^ (Walker A) or _PARL_Skd3^Y430A^ (pore loop). _PARL_Skd3^K387A^ (Walker A) subunits inhibited _PARL_Skd3 ATPase activity in a manner that suggested the incorporation of ∼3-5 mutant subunits inactivate the hexamer (Figure 6E). Thus, Skd3 ATPase activity appears to be sub-globally cooperative. By contrast, titrating _PARL_Skd3^Y430A^ (pore loop) subunits did not affect _PARL_Skd3 ATPase activity any more than dilution in buffer (Figure 6F). Hence, the ATPase activity of the _PARL_Skd3 hexamer is more resistant to pore-loop mutant subunits than Walker A mutant subunits. These findings contrast with observations made with Hsp104 (DeSantis et al. 2012) where Walker A mutant subunits do not affect the ATPase activity of the hexamer more than dilution in buffer and pore-loop mutant subunits have no effect (DeSantis et al. 2012). Thus, Hsp104 and _PARL_Skd3 appear to display different subunit co-operativity with respect to ATP hydrolysis.

We next examined how _PARL_Skd3^K387A^ (Walker A) and _PARL_Skd3^Y430A^ (pore loop) subunits affected _PARL_Skd3 disaggregase activity. Incorporation of two _PARL_Skd3^K387A^ (Walker A) subunits is sufficient to inactivate the _PARL_Skd3 hexamer (Figure 6G). Thus, _PARL_Skd3 hexamers are very sensitive to individual subunits that are not able to bind or hydrolyze ATP due to a defective Walker A motif. This finding reinforces our earlier observation that _PARL_Skd3 disaggregase activity is sharply inhibited by the slowly hydrolyzable ATP analog, ATPγS (Figure 5E, F). By contrast, incorporation of five _PARL_Skd3^Y430A^ (pore loop) subunits is needed to inactivate the _PARL_Skd3 hexamer (Figure 6H). Thus, even though _PARL_Skd3^Y430A^ has reduced ATPase activity (Cupo and Shorter, 2020b), it appears that this ATPase defect is more readily buffered by WT subunits upon incorporation into _PARL_Skd3 hexamers. It is also likely that substrate release caused by insufficient binding of the pore loops contributes to inhibition by _PARL_Skd3^Y430A^ subunits. Overall, these findings suggest that _PARL_Skd3 utilizes a subglobally cooperative mechanism to disaggregate disordered luciferase aggregates. Thus, at least five subunits must have a functional Walker A motif to bind and hydrolyze ATP and at least two subunits must be able to engage substrate tightly via Y430 for productive luciferase disaggregase activity. Viewed in another way, our findings illustrate that _PARL_Skd3 hexamers display robustness and can buffer a specific number of mutant subunits. Indeed, _PARL_Skd3 hexamers can tolerate one subunit with a defective Walker A motif unable to bind and hydrolyze ATP, and four subunits with a defective pore loop and still drive luciferase disaggregation. This robustness has implications for the etiology of SCN and MGCA7.

### SCN-linked subunits inhibit _PARL_Skd3 activity more severely than MGCA7-linked **_PARL_Skd3 subunits**

Next, we surveyed the location of disease-linked mutations in the Skd3 structure. Biallelic MGCA7-linked mutations are scattered throughout Skd3 (Figure 7A) (Wortmann et al., 2015) MGCA7-linked mutations have been found in the MTS, ANK, NBD, and CTD (Figure 7A). For example, T268M, A269T, and Y272C cluster in the third ankyrin repeat of the ANK and are highly conserved residues of the ankyrin-repeat motif (Figure 7A). Specifically, residues T268 and A269 lie near the N-terminal portion of the first alpha helix in the third ankyrin repeat (Figure S3F). Mutations at either residue would be predicted to destabilize the alpha helix and potentially disrupt the fold of the entire third ankyrin repeat (Mosavi et al., 2002). Other MGCA7-linked mutations are found in the large (e.g. M411I, R460P, C486R, and E501K) and small (e.g. A591V, R628C, and R650P) subdomains of the NBD (Figure 7A). Some MGCA7-linked NBD mutations such as R475Q and R408G are in residues that form interprotomer contacts (Figure 7B).

**Figure 7.**
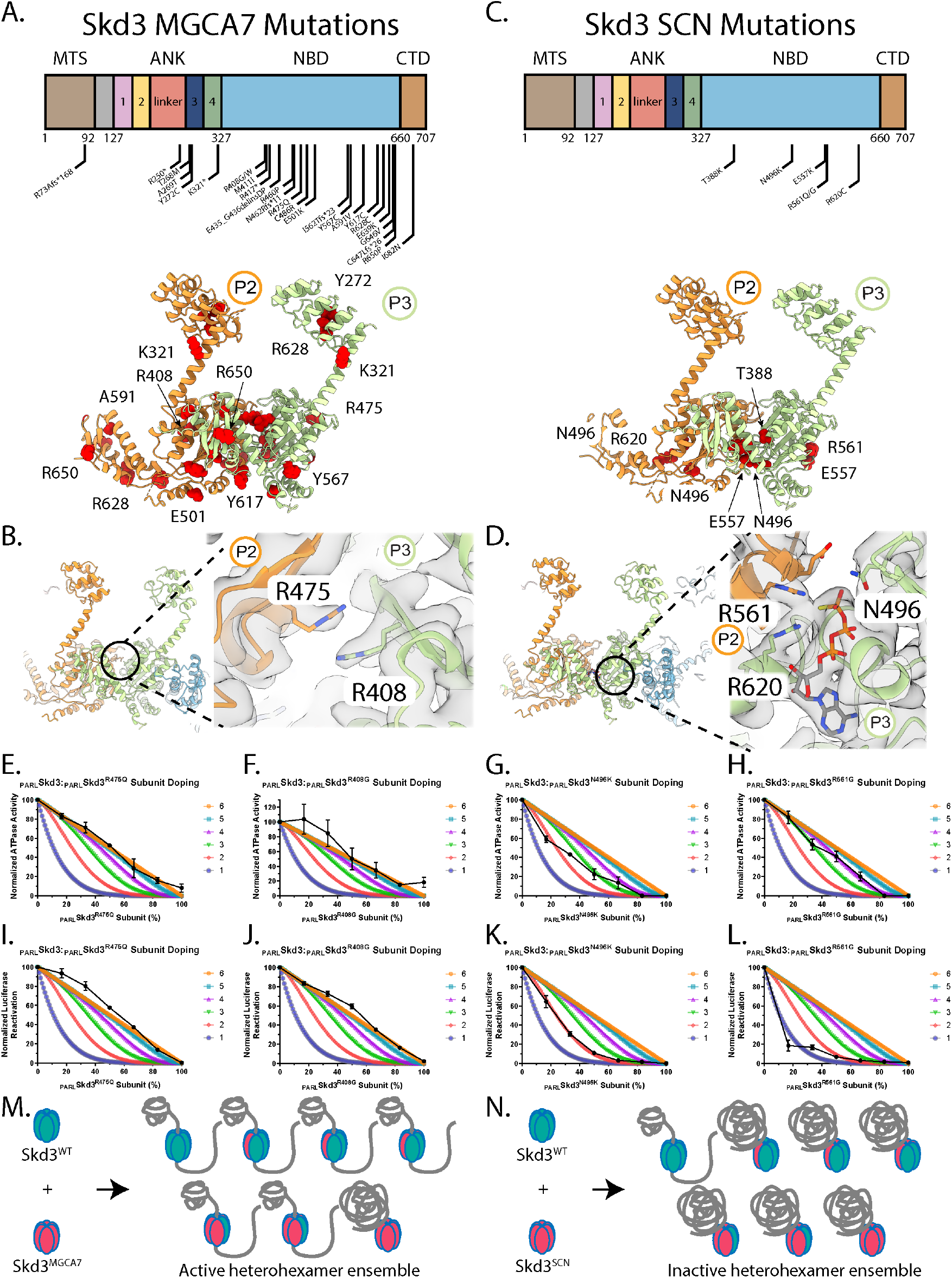
SCN-linked subunits inhibit _PARL_Skd3 activity more severely than MGCA7-linked _PARL_Skd3 subunits. **(A)** Domain map depicting all published biallelic mutations in Skd3 that have been associated with MGCA7 (top). Model of protomers P2 and P3 with MGCA7-linked mutations colored in red (bottom). **(B)** Model of back protomers colored by individual protomers (left). Interaction interface of residue R475 from protomer P2 and residue R408 from protomer P3. **(C)** Domain map depicting all published mutations in Skd3 that have been associated with SCN. Model of protomers P2 and P3 with SCN-linked mutations colored in red. **(D)** Model of back protomers colored by individual protomers (left). Interaction interface of residues E557 and R561 of protomer P2 and residues N496 and R620 from protomer P3 within the nucleotide binding pocket of protomer P3. **(E-H)** ATPase activity of _PARL_Skd3 mixed with various ratios of _PARL_Skd3^R475Q^ (E), _PARL_Skd3^R408G^ (F), _PARL_Skd3^N496K^ (G), or _PARL_Skd3^R561G^ (H). ATPase activity was buffer subtracted and normalized to _PARL_Skd3 (N = 3, data shown as black dots with mean ± SEM). **(I-L)** Luciferase disaggregase activity of _PARL_Skd3 mixed with various ratios of _PARL_Skd3^R475Q^ (I), _PARL_Skd3^R408G^ (J), _PARL_Skd3^N496K^ (K), or _PARL_Skd3^R561G^ (L). Disaggregase activity was buffer subtracted and normalized to _PARL_Skd3 (N = 3, data shown as black dots with mean ± SEM). **(M)** Schematic illustrating that _PARL_Skd3 hexamers containing a mixture of WT and MGCA7-linked subunits are typically active disaggregases. **(N)** Schematic illustrating that _PARL_Skd3 hexamers containing a mixture of WT and SCN-linked subunits are typically less active disaggregases. See also Figure S6.

By contrast, SCN-linked mutations have only been identified in the NBD and cluster specifically within the nucleotide-binding pocket (Figure 7C, D) (Warren et al., 2022). Remarkably, most of the SCN-linked mutations are in canonical AAA+ motifs. For example, N496K mutates the sensor-1 motif, which coordinates the attacking water molecule relative to the γ-phosphate of ATP and transmits a conformational change upon nucleotide engagement to displace the Arg finger in the adjacent nucleotide pocket (Figure 2E, S2I, 7D) (Erzberger and Berger, 2006; Hanson and Whiteheart, 2005; Puchades et al., 2020; Wendler et al., 2012). R561G mutates the Arg-finger residue, which contacts the γ-phosphate of ATP in the nucleotide-binding pocket of the adjacent protomer and is key for ATP hydrolysis (Figure 2E, S2I, 7D) (Erzberger and Berger, 2006; Hanson and Whiteheart, 2005; Puchades et al., 2020; Wendler et al., 2012). R620C mutates the sensor-2 motif, which contacts both the β- and γ-phosphate of ATP to mediate a conformational change that sequesters the catalytic site from water (Figure 2E, S2I, 7D) (Erzberger and Berger, 2006; Hanson and Whiteheart, 2005; Puchades et al., 2020; Wendler et al., 2012). T388K is directly adjacent to the Walker A motif, is highly conserved among other HCLR clade AAA+ proteins, and faces into the nucleotide-binding pocket (Cupo and Shorter, 2020b) (Figure 2E, 7C). Similarly, E557K lies within a conserved stretch of residues near the Arg finger and also makes contact with nucleotide (Figure 7C, D) (Cupo and Shorter, 2020b).

MGCA7-linked mutations impair disaggregase activity in a manner that predicts disease severity (Cupo and Shorter, 2020b). However, MGCA7-linked mutations do not always impair ATPase activity (Cupo and Shorter, 2020b). By contrast, SCN-linked mutations impair ATPase and disaggregase activity (Warren et al., 2022). It is not understood why SCN-linked mutations are dominant-negative, whereas MGCA7-linked mutations are recessive. To assess how severely MGCA7-linked and SCN-linked variants affect WT Skd3 activity, we selected three variants associated with each disease for subunit doping studies. Specifically, we used MGCA7-linked variants: _PARL_Skd3^R408G^, _PARL_Skd3^R475Q^, and _PARL_Skd3^A591V^ (Figure 7A, B), and SCN-linked variants: _PARL_Skd3^N496K^, _PARL_Skd3^R561G^, and _PARL_Skd3^R620C^ (Figure 7C, D) (Pronicka et al., 2017; Warren et al., 2022). All of these disease-linked Skd3 variants are severely impaired for disaggregase activity (Cupo and Shorter, 2020b; Warren et al., 2022). Likewise, these disease-linked variants all have diminished ATPase activity, with the exception of the MGCA7-linked variant _PARL_Skd3^R408G^, which exhibits ∼20% of WT ATPase activity (Cupo and Shorter, 2020b; Warren et al., 2022).

We assessed the ability of the disease-linked _PARL_Skd3 variants to form hexamers and dodecamers in the presence of ATPγS and absence of substrate. Unlike _PARL_Skd3, which was shifted toward the hexameric form, two of the MGCA7-linked variants, _PARL_Skd3^R408G^ and _PARL_Skd3^R475Q^, were shifted toward dodecamers (Figure S6A). By contrast, MGCA7-linked _PARL_Skd3^A591V^ was shifted to lower molecular weight oligomers (Figure S6A). _PARL_Skd3^A591V^ formed some hexamers, but the dodecameric species was reduced (Figure S6A). The SCN-linked variant, _PARL_Skd3^N496K^, was shifted toward the hexameric form like _PARL_Skd3 (Figure S6B). The remaining SCN-linked variants, _PARL_Skd3^R561G^ and _PARL_Skd3^R620C^, were shifted toward the dodecameric form (Figure S6B).

To test how severely each disease-linked variant affected WT _PARL_Skd3 activity, we mixed each disease-linked variant and WT _PARL_Skd3 and assessed how they affected ATPase activity. Importantly, FRET studies revealed that disease-linked _PARL_Skd3 subunits were effectively incorporated into WT _PARL_Skd3 hexamers (Figure S6C, D). Addition of the MGCA7-linked variant _PARL_Skd3^R475Q^ to _PARL_Skd3 revealed that six _PARL_Skd3^R475Q^ subunits are necessary to reduce ATPase activity to the same level as _PARL_Skd3^R475Q^ (Figure 7E). _PARL_Skd3^R408G^ retains ∼20% of WT ATPase activity and five _PARL_Skd3^R408G^ subunits per hexamer reduced ATPase activity to this level (Figure 7F). Likewise, five _PARL_Skd3^A591V^ subunits per hexamer are required to eliminate ATPase activity (Figure S6E). These findings suggest that MGCA7-linked subunits have very mild effects on the ATPase activity of WT subunits.

The inhibitory effect of SCN-linked subunits on ATPase activity was more pronounced than that of MGCA7-liked subunits. Indeed, all SCN-linked variants more effectively inhibited the ATPase activity of _PARL_Skd3 in subunit mixing experiments (Figure 7G, H, S6G). Incorporation of two to four _PARL_Skd3^N496K^ subunits inactivated the hexamer (Figure 7G). Moreover, three to four SCN-linked _PARL_Skd3^R561G^ subunits or four _PARL_Skd3^R620C^ subunits inactivated the hexamer (Figure 7H, S6G). Thus, SCN-linked subunits more sharply inhibit the ATPase activity of WT subunits than MGCA7-linked subunits.

Next, we assessed how MGCA7-linked subunits affected _PARL_Skd3 disaggregase activity in mixing experiments. Six MGCA7-linked _PARL_Skd3^R408G^, _PARL_Skd3^R475Q^, or _PARL_Skd3^A591V^ subunits were needed to eliminate _PARL_Skd3 disaggregase activity (Figure 7I, J, S6F). Strikingly, _PARL_Skd3^A591V^ subunits barely affected disaggregase activity even when 3 mutant subunits were incorporated into the hexamer (Figure S6F). Thus, even one WT _PARL_Skd3 subunit in an otherwise _PARL_Skd3^R408G^, _PARL_Skd3^R475Q^, or _PARL_Skd3^A591V^ hexamer enables disaggregase activity. These findings suggest that MGCA7-linked mutant subunits typically have only minor effects on the disaggregase activity of WT subunits within the hexamer. The strong buffering activity of WT _PARL_Skd3 subunits provides a mechanistic explanation for why MGCA7-linked mutations are biallelic and recessive.

Finally, we assessed how SCN-linked subunits affected _PARL_Skd3 disaggregase activity in mixing experiments (Figure 7K, L, S6H). Incorporation of two SCN-linked _PARL_Skd3^N496K^ subunits was sufficient to inactivate the hexamer (Figure 7K). SCN-linked _PARL_Skd3^R561G^ subunits had the most drastic inhibitory effects. Only one or two _PARL_Skd3^R561G^ subunits were required to inactivate the hexamer (Figure 7L). Finally, incorporation of three SCN-linked _PARL_Skd3^R620C^ subunits inactivated the hexamer (Figure S6H). In sum, our findings strongly suggest that _PARL_Skd3 utilizes a subglobally co-operative mechanism to disaggregate luciferase. Moreover, SCN-linked subunits generally have a sharper inhibitory effect on WT _PARL_Skd3 than MGCA7-linked subunits (Figure 7E-L, S6E-J). These results provide a mechanistic explanation for why SCN-linked mutations are dominant negative and typically monoallelic.

## Discussion

Here, we describe the first structures of _PARL_Skd3 (human *CLPB*) and define mechanisms by which _PARL_Skd3 drives protein disaggregation. _PARL_Skd3 forms a hexameric complex with an asymmetric seam between protomers P1 and P6, analogous to other AAA+ proteins such as Hsp104, ClpB, and ClpA (Gates et al., 2017; Lopez et al., 2020; Rizo et al., 2019). _PARL_Skd3 subunits adopt a hexameric arrangement that engages substrate in its central channel via pore-loop interactions in the NBD. Indeed, _PARL_Skd3 likely employs a conserved translocation mechanism identified in other AAA+ disaggregases and translocases (Gates et al., 2017; Puchades et al., 2017; Rizo et al., 2019). Mutation of conserved primary pore-loop residues that engage substrate (e.g. Y430 and V431) reduce protein disaggregase activity (Cupo and Shorter, 2020b). Interestingly, mutations at V431 are observed in the human population (V431D, V431A, and V431I) with low frequency according to the Genome Aggregation Database, although none of the known carriers are homozygous (Karczewski et al., 2020). Based on our data, we predict that specific biallelic mutations to V431 would be highly pathogenic.

_PARL_Skd3 also contains a secondary pore loop that engages substrate. An R417A mutation ablated _PARL_Skd3 disaggregase activity but had no effect on ATPase activity, indicating a critical role for this arginine. By contrast, mutations in the secondary pore loops of Hsp104 and ClpB have much milder effects on disaggregase activity (Howard et al., 2020; Rizo et al., 2019). Thus, the secondary pore loop of _PARL_Skd3 plays a more important role in disaggregase activity. We suggest that the guanidyl group of the R417 side chain may create a local denaturing microenvironment, which maintains the unfolded state of the polypeptide as it is extruded from the _PARL_Skd3 channel. Indeed, the six R417 residues facing into the central _PARL_Skd3 channel create a local guanidine concentration of ∼11.6 M. In this way, R417 might serve as an ‘arginine denaturation collar’ akin to those proposed for other AAA+ proteins such as p97/VCP and Vps4 (DeLaBarre et al., 2006; Gonciarz et al., 2008).

One of the most prominent and unique features of the _PARL_Skd3 structure is the presence of a dodecameric species, created by two hexamers making head-to-head contacts through the ANK domain. The hexamer and dodecamer exist in dynamic equilibrium, but the dodecamer predominates upon polypeptide binding. The head-to-head ANK contacts could concentrate _PARL_Skd3 disaggregases on the aggregate surface and enable stronger pulling forces by maximizing the number of hexamers simultaneously processing substrate at once. Indeed, _PARL_Skd3^ΔL^, which lacks the linker region in the ANK, exhibited reduced dodecamer formation and reduced disaggregase activity, whereas ATPase activity was unaffected. Moreover, _PARL_Skd3^ΔL507-I534^, which lacks the novel insertion in the NBD, exhibits increased dodecamer formation, disaggregase activity, and ATPase activity. Our findings suggest that dodecamer formation enhances _PARL_Skd3 disaggregase activity.

The ankyrin repeats are another unique feature of the _PARL_Skd3 structure. To the best of our knowledge, Skd3 is the only protein that combines a AAA+ domain with ankyrin repeats. The ANK and NBD are required for Skd3 ATPase and disaggregase activity (Cupo and Shorter, 2020b). Alpha fold predicts that N-terminal ankyrin repeats 1 and 2 stack on the C-terminal ankyrin repeats 3 and 4, with the largely disordered linker excluded (Figure 3A). Deletion of ankyrin repeats 1 and 2, the linker, or ankyrin repeats 3 and 4 from the ANK reduces disaggregase activity, but not ATPase activity. However, each of these deletion variants retained ∼20-45% _PARL_Skd3 disaggregase activity, indicating that the remaining ankyrin repeats and linker can support some activity. The linker region promotes dodecamer formation, whereas deletion of ankyrin repeats 1 and 2 or 3 and 4 does not perturb dodecamerization. The ankyrin repeats could play a role in substrate engagement or disaggregase plasticity analogous to the Hsp104 N-terminal domain (Sweeny et al., 2015; Sweeny et al., 2020; Wang et al., 2017).

Interestingly, several Skd3 transcript variants are present in humans, which differ only within the ANK (The UniProt Consortium, 2021). Residues R152-N180, corresponding to part of ankyrin-repeats 1 and 2, are absent in transcript variant 3. Residues D216-G245, corresponding to the middle section of the linker, are absent in transcript variants 2 and 3. The functional consequences of these transcript variants are not clear, but both deletions correspond to the length of almost exactly one ankyrin repeat (Figure 1A). We suggest that cells may tune the level of Skd3 disaggregase activity via translation of these distinct Skd3 transcripts.

Skd3 also contains a 28 amino acid insertion in the NBD between the sensor-1 and Arg-finger motifs (Cupo and Shorter, 2020b). Alpha fold predicts that this stretch of residues extends as a helix protruding from the NBD. Deleting this helix results in enhanced dodecamer formation and accelerated ATPase activity and disaggregase activity. We propose that this insertion acts as a regulatory element to slow the ATPase motor. Skd3 also has an extended CTD that is patterned with both acidic and basic residues (Cupo and Shorter, 2020b). This patterning contrasts with the acidic extended CTD of Hsp104 (Mackay et al., 2008). Deleting the CTD slightly accelerates _PARL_Skd3 ATPase but not disaggregase activity. Thus, the CTD appears to enable efficient coupling of ATP hydrolysis and mechanical work.

Hsp104 operates at low millimolar concentrations of ATP and is potently inhibited by ADP (Grimminger et al., 2004; Klosowska et al., 2016). By contrast, _PARL_Skd3 can operate at low micromolar concentrations of ATP and is less potently inhibited by ADP. _PARL_Skd3 is likely adapted to lower concentrations of ATP and lower ratios of ATP:ADP found within mitochondria (Heldt et al., 1972). Under stressed conditions where mitochondrial function is impaired, the ratio of ATP:ADP will decrease further. Thus, to preserve mitochondrial fitness _PARL_Skd3 must remain functional. In principle, _PARL_Skd3 could act as a determinant of cell fate whereby _PARL_Skd3 preserves mitochondrial function until a critical ratio of ATP:ADP has been breached. After this point, _PARL_Skd3 would no longer chaperone the mitochondrial intermembrane space to maintain cell viability.

_PARL_Skd3 disaggregase activity was very sensitive to slowly hydrolyzable ATPγS. Even an 11:1 ratio of ATP: ATPγS strongly inhibited _PARL_Skd3 disaggregase activity. Thus, _PARL_Skd3 disaggregase activity is sensitive to individual subunits that hydrolyze ATP slowly. Indeed, our mutant subunit-doping studies suggest that _PARL_Skd3 utilizes a subglobally cooperative mechanism (i.e. 2-5 subunits collaborate) to disaggregate disordered luciferase aggregates. _PARL_Skd3 disaggregase activity was very sensitive to mutant subunits that were defective in ATP hydrolysis. Thus, one or two Arg-finger mutant (R561G) subunits, two Walker A mutant (K387A) or sensor-1 mutant (N496K) subunits, or three sensor-2 mutant (R620C) subunits per _PARL_Skd3 hexamer ablated activity. _PARL_Skd3 hexamers exhibit some robustness and can buffer the incorporation of a specific number of mutant subunits, i.e. one subunit with a defective Walker-A motif or sensor-1 motif and two subunits with a defective sensor-2 motif. These results reveal that some AAA+ motifs are likely more important for subunit co-operativity within the hexamer than others. For example, the Arg finger appears to be more critical than the sensor-2 motif. _PARL_Skd3 hexamers also exhibited robustness against subunits with a defective primary pore loop (Y430A). Thus, _PARL_Skd3 could tolerate four subunits with the Y430A mutation, indicating that two functional pore loops are required for _PARL_Skd3 to maintain a grip on substrate during disaggregation. Overall, these findings differ from prior observations with Hsp104, which uses a probabilistic mechanism to disaggregate disordered luciferase aggregates (DeSantis et al., 2012).

Recently, Skd3 has been highlighted as a potential therapeutic target for the treatment of several cancers (Chen et al., 2019; Pudova et al., 2020). Our structures of _PARL_Skd3 will enable computational drug design and drug discovery for small-molecule inhibitors of Skd3. They are also useful for interpreting mutations linked to MGCA7 and SCN. Indeed, SCN-linked mutations cluster within the nucleotide-binding pocket, whereas MGCA7-linked mutations are scattered throughout Skd3, including in the third ankyrin repeat of the ANK, at the protomer-protomer interface of the NBD, and within the small domain of the NBD. Importantly, we establish that SCN-linked mutant subunits more sharply inhibit _PARL_Skd3 ATPase and disaggregase activity than MGCA7-linked subunits. The robustness of _PARL_Skd3 against inhibition by MGCA7-linked subunits provides a mechanistic explanation for why MGCA7-linked mutations are recessive and must be biallelic to cause disease. Moreover, the sharp inhibition by SCN-linked mutant subunits provides a mechanistic explanation for why SCN-linked mutations are dominant negative.

Both MGCA7 and SCN are characterized by loss-of-function Skd3 mutations (Cupo and Shorter, 2020b; Kanabus et al., 2015; Kiykim et al., 2016; Pronicka et al., 2017; Saunders et al., 2015; Warren et al., 2022; Wortmann et al., 2016; Wortmann et al., 2021; Wortmann et al., 2015). Thus, in principle, both diseases could be treated by increasing Skd3 activity. However, due to the mechanistic differences between how MGCA7-linked and SCN-linked subunits affect the activity of WT _PARL_Skd3, different treatment modalities will likely be beneficial for each disease. For treating biallelic MGCA7 mutations, expression of WT Skd3 via adeno-associated viruses (AAV) is a viable therapeutic option (Kuzmin et al., 2021). Indeed, expression of WT genes via AAV has yielded FDA-approved therapies for congenital blindness and spinal muscular atrophy (Al-Zaidy et al., 2019; Apte, 2018; Mendell et al., 2017). Here, the robustness of _PARL_Skd3 hexamers against MGCA7-linked subunits will enable restoration of _PARL_Skd3 activity (Figure 7M, S6I). By contrast, SCN-linked mutations are dominant negative, and SCN-linked subunits sharply inhibit WT _PARL_Skd3. Consequently, an AAV strategy to deliver the WT Skd3 gene is likely to be less effective for SCN (Figure 7N, S6J). We suggest that a therapeutic strategy that reduces or eliminates expression of the mutant Skd3 allele is likely to be more beneficial for SCN. Here, gene editing, specific antisense oligonucleotides, or AAV-delivered siRNA to specifically reduce mutant allele expression could be viable therapeutic strategies to enable restoration of Skd3 activity (Crooke et al., 2021; Frangoul et al., 2021; Kuzmin et al., 2021; Malech, 2021).

## Experimental Procedures

### Multiple sequence alignments

NBD sequences were acquired via UniProtKB for Homo sapiens Skd3, Escherichia coli ClpA, Escherichia coli ClpB, Staphylococcus aureus ClpC, Escherichia coli ClpX, Saccharomyces cerevisiae Hsp78, Arabidopsis thaliana Hsp101, Saccharomyces cerevisiae Hsp104, Escherichia coli RuvB, and Pseudomonas aeruginosa ClpG. Ankyrin-repeat sequences were acquired via UniProtKB for Homo sapiens Skd3. Consensus ankyrin repeat was derived from Mosavi, et. al. (Mosavi et al., 2002). Compiled sequences were aligned via Clustal Omega (Madeira et al., 2019). The linker region of the ankyrin repeats was aligned manually to the Clustal Omega alignment. Alignment image was generated via BoxShade tool as described previously (Cupo and Shorter, 2020b).

### Purification of _PARL_Skd3

_PARL_Skd3 and variants were purified as previously described (Cupo and Shorter, 2020a, b). In short, _PARL_Skd3 and variants were expressed with an N-terminal MBP-tag in BL21 (DE3) RIL cells (Agilent). Cells were lysed via sonication in lysis buffer (40 mM HEPES-KOH pH = 7.4, 500 mM KCl, 20% [w/v] glycerol, 5 mM ATP, 10 mM MgCl_2_, 2 mM β-mercaptoethanol, 2.5 µM PepstatinA, and cOmplete Protease Inhibitor Cocktail [one tablet/250 mL, Millipore Sigma]). Lysates were cleared via centrifugation at 30,597xg and 4°C for 20 min and the supernatant was applied to amylose resin (NEB). The column was washed with 15 column volumes (CV) of wash buffer (WB: 40 mM HEPES-KOH pH = 7.4, 500 mM KCl, 20% [w/v] glycerol, 5 mM ATP, 10 mM MgCl_2_, 2 mM β-mercaptoethanol, 2.5 µM PepstatinA, and cOmplete Protease Inhibitor Cocktail [1 full size tablet/50mL, Millipore Sigma]) at 4°C, 3 CV of WB supplemented with 20 mM ATP at 25°C for 30 min, and an additional 15 CV of WB at 4°C. The protein was then washed with ∼8 CV of elution buffer (EB: 50 mM Tris-HCl pH = 8.0, 300 mM KCl, 10% glycerol, 5 mM ATP, 10 mM MgCl_2_, and 2 mM β-mercaptoethanol) and eluted via TEV protease cleavage at 34°C. The protein was run over a size exclusion column (GE Healthcare HiPrep 26/60 Sephacryl S-300 HR) in sizing buffer (50 mM Tris-HCl pH = 8.0, 500 mM KCl, 10% glycerol, 1 mM ATP, 10 mM MgCl_2_, and 1 mM DTT). Peak fractions were collected, concentrated to ∼5 mg/mL, supplemented with 5 mM ATP, and snap frozen. Protein purity was determined to be >95% by SDS-PAGE and Coomassie staining.

### Purification of Hsp104

Hsp104 was purified as previously described (DeSantis et al., 2012). In short, Hsp104 was expressed in BL21 (DE3) RIL cells, lysed via sonication in lysis buffer (50 mM Tris-HCl pH = 8.0, 10 mM MgCl_2_, 2.5% glycerol, 2 mM β-mercaptoethanol, 2.5 µM PepstatinA, and cOmplete Protease Inhibitor Cocktail [one mini EDTA-free tablet/50 mL, Millipore Sigma]), centrifuged at 30,597xg and 4°C for 20 min, and purified on Affi-Gel Blue Gel (Bio-Rad). Hsp104 was eluted in elution buffer (50 mM Tris-HCl pH = 8.0, 1M KCl, 10 mM MgCl_2_, 2.5% glycerol, and 2 mM β-mercaptoethanol) and then exchanged into storage buffer (40 mM HEPES-KOH pH = 7.4, 500 mM KCl, 20 mM MgCl2, 10% glycerol, 1 mM DTT). The protein was diluted to 10% in buffer Q (20 mM Tris-HCl pH = 8.0, 50 mM NaCl, 5 mM MgCl_2_, and 0.5 mM EDTA) and loaded onto a 5 mL RESOURCE Q anion exchange chromatography (GE Healthcare). Hsp104 was eluted via linear gradient of buffer Q+ (20 mM Tris pH = 8.0, 1M NaCl, 5 mM MgCl_2_, and 0.5 mM EDTA). The protein was exchanged into storage buffer and snap frozen. Protein purity was determined to be >95% by SDS-PAGE and Coomassie staining.

### Purification of Hsc70 and Hdj1

Hsc70 and Hdj1 were purified as previously described (Michalska et al., 2019). Hsc70 and Hdj1 were expressed in BL21 (DE3) RIL cells with an N-terminal His-SUMO tag. Cells were lysed via sonication into lysis buffer (50 mM HEPES-KOH pH = 7.5, 750 mM KCl, 5 mM MgCl_2_, 10% glycerol, 20 mM imidazole, 2 mM β-mercaptoethanol, 5 µM pepstatin A, and cOmplete Protease Inhibitor Cocktail [one mini EDTA-free tablet/50 mL, Millipore Sigma]). Lysates were cleared via centrifugation at 30,597xg and 4°C for 20 min. The supernatant was bound to Ni-NTA Agarose resin (Qiagen), washed with 10 CV of wash buffer (50 mM HEPES-KOH pH = 7.5, 750 mM KCl, 10 mM MgCl_2_, 10% glycerol, 20 mM imidazole, 1 mM ATP, and 2 mM β-mercaptoethanol), and eluted with 2 CV of elution buffer (wash buffer supplemented with 300 mM imidazole). The tag was removed via Ulp1 (1:100 Ulp1:Protein molar ratio) cleavage during dialysis into wash buffer. The protein was further purified via loading onto a 5 mL HisTrap HP column (GE Healthcare) and pooling the untagged elution. The protein was pooled and concentrated, and then purified further via Resource Q ion exchange chromatography. The elution was pooled, concentrated, and snap frozen. Protein purity was determined to be >95% via SDS-PAGE and Coomassie staining.

### Size-exclusion chromatography

All size-exclusion chromatography experiments were run on a Superose 6 Increase 3.2/300 (Cytiva) column pre-equilibrated in buffer containing: 40 mM HEPES (pH = 8.0), 40 mM KCl, 10 MgCl_2_, and 1 mM DTT. To form a substrate-bound complex, _PARL_Skd3 (20 µM) was incubated with FITC-casein (55 µM) (#C0528; Sigma) in the presence of nucleotide (ATPγS, ATP, ADP, or AMP-PNP) (5 mM) for 15 min at room temperature. For experiments without FITC-casein, _PARL_Skd3 (20 µM) and nucleotide (ATPγS, ATP, ADP, or AMP-PNP) (5 mM) were incubated for 15 minutes at room temperature. After the incubation period, the samples were spin-filtered before injecting on column.

### Cryo-EM Data Collection and Processing for _PARL_Skd3:casein:ATPγS Complex

To form a substrate-bound complex, _PARL_Skd3 (55 µM) was incubated with FITC-casein (55 µM) (#C0528; Sigma) in the presence of ATPγS (5 mM) in buffer containing: 40 mM HEPES (pH = 8.0), 40 mM KCl, 10 MgCl_2_, 1 mM DTT. After incubating for 15 minutes at room temperature, the sample was applied to a Superose 6 Increase 3.2/300 column (GE Healthcare) for size exclusion chromatography (SEC) analysis. The fraction corresponding to the largest molecular weight complex from SEC of _PARL_Skd3 and FITC-casein (Figure 1A) was isolated and incubated with 1 mM ATPγS. Before freezing, proper dilutions were made to a final concentration of ∼.7 mg/mL and a 3.0 µL drop was applied to glow discharged holey carbon (R 1.2/1.3; Quantifoil), then blotted for 3 s. at 4°C and 100% humidity with a blot force of 1, followed by an additional 3.0µL drop. The sample was then blotted again for 2 s. with a blot force of 0 with Whatman No. 1 filter paper before being plunge frozen in liquid ethane using a Vitrobot (Thermo Fischer Scientific).

The sample was then imaged on a Titan Krios TEM (Thermo Fischer Scientific) operated at 300 keV and equipped with a Gatan BioQuantum imaging energy filter using a 20eV zero loss energy slit (Gatan Inc). Movies were acquired in super-resolution mode on a K3 direct electron detector (Gatan Inc.) at a calibrated magnification of 58,600X corresponding to a pixel size of 0.4265 Å/pixel. A defocus range of 0.8 to 1.2 μm was used with a total exposure time of 2 seconds fractionated into 0.2s subframes for a total dose of 68 e-/Å^2^ at a dose rate of 25 e- /pixel/s. Movies were subsequently corrected for drift using MotionCor2 (Zheng et al., 2017) and were Fourier-cropped by a factor of 2 to a final pixel size of 0.853 Å/pixel.

A total of ∼30,000 micrographs were collected over multiple datasets. Micrograph quality was assessed and poor micrographs, including those above the resolution cutoff of ∼5Å, were discarded. The individual datasets were processed separately to ensure data quality before combining them all together for further processing. Data processing was performed in cryoSPARC v3.2 (Punjani et al., 2017). For particle picking, blob picker was set to 180 Å-200 Å for minimum and maximum particle diameter and the particles picked were inspected before extracting particles. 2D classification was performed to remove contamination and junk particles and good classes were selected which left ∼900,000 remaining particles. Four different ab-initio models were reconstructed which were then used in 3D classification.

Heterogenous refinement was performed with 4 different classes, which resulted in 3 distinct classes: hexamer, Class 1 (41%, ∼358K particles); heptamer, Class 3 (19%, ∼167K particles); and dodecamer, Class 2 (15%, ∼130K particles); Other, Class 4 (24%, ∼210K particles) (Figure S1B). Each of the classes underwent homogenous refinement which resulted in resolutions of 9Å for the dodecamer, 7 Å for the heptamer and 3.2Å for the hexamer. To improve the resolution, non-uniform refinement was completed on both the dodecamer and hexamer to improve the resolutions to 7.2Å and 2.9Å respectively (Figure S1D and S3B). Both classes underwent local CTF refinement which did not result in an improvement in resolution.

### Cryo-EM Data Collection and Processing for _PARL_Skd3^Δ1-2^:casein:ATPγS Complex

To form a substrate-bound complex, _PARL_Skd3^Δ1-2^ (40 µM) was incubated with FITC-casein (40 µM) (#C0528; Sigma) in the presence of ATPγS (5 mM) in buffer containing: 40 mM HEPES (pH = 8.0), 40 mM KCl, 10 MgCl_2_, 1 mM DTT. After incubating for 15 minutes at room temperature, grids were prepared. For grid freezing, a 3.0 µL drop was applied to glow discharged holey carbon (R 1.2/1.3; Quantifoil), then blotted for 3 s. at 4°C and 100% humidity with a blot force of 1 followed by an additional 3.0µL drop. The sample was then blotted again for 2 s. with a blot force of 0 with Whatman No. 1 filter paper before being plunge frozen in liquid ethane using a Vitrobot (Thermo Fischer Scientific).

The sample was then imaged on a Glacios TEM (Thermo Fischer Scientific) operated at 200 keV (Gatan Inc). Movies were acquired in super-resolution mode on a K2 direct electron detector (Gatan Inc.) at a calibrated magnification of 108,695X corresponding to a pixel size of 0.463 Å/pixel. A defocus range of 1.0 to 2.0 μm was used with a total exposure time of 6 seconds fractionated into 0.06s subframes for a total dose of 55.8 e-/Å^2^ at a dose rate of 8 e-/pixel/s. Movies were subsequently corrected for drift using MotionCor2 (Zheng et al., 2017) and were Fourier-cropped by a factor of 2 to a final pixel size of 0.972 Å/pixel.

A total of ∼15,000 micrographs were collected over multiple datasets. The individual datasets were processed separately to ensure data quality before combining them all together for further processing. Micrograph quality was assessed and poor micrographs, including those above the resolution cutoff of ∼5Å, were discarded. Data processing was performed in cryoSPARC v3.2 (Punjani et al., 2017). For particle picking, blob picker was set to 180 Å-200 Å for minimum and maximum particle diameter and the particles picked were inspected before extracting particles. 2D classification was performed in two rounds to remove contamination and junk particles and good classes were selected which left ∼700,000 remaining particles. The results from a previous 3D classification were used as the starting models for 3D classification.

Heterogenous refinement was performed with 4 different classes, which resulted in 3 distinct classes: dodecamer, Class 1 (24%, ∼165K particles); bent dodecamer, Class 2 (30%, ∼203K particles); trimer, Class 3 (32%, ∼213K particles); and other, Class 4 (14%, ∼95K particles) (Figure S3I). Each of the classes underwent homogenous refinement which resulted in resolutions of 8Å for the dodecamer, 7Å for the bent dodecamer and 8Å for the trimer (Figure S3I-M).

### Molecular Modeling

An initial model for _PARL_Skd3was generated in SWISS-MODEL (Waterhouse et al., 2018) and was docked into the EM map using the UCSF chimera’s function fit in map (Pettersen et al., 2004). The initial model lacked the ANK so the SWISS-MODEL generated was combined with the Alpha-fold prediction of the ANK taken from the AlphaFold Protein Structure Database. Initial refinement was performed using Rosetta_Relax in cartesian space to generate 30 different models. The map/model quality for each model generated was examined in Chimera (Pettersen et al., 2004) and the lowest energy minimized model was used moving forward. Various outliers and poorly fit density were manually fixed using ISOLDE (Croll, 2018) in ChimeraX (Pettersen et al., 2021). To fix most of the outliers another round of Rosetta_Relax in cartesian space was performed followed by iterative rounds of refinement in Phenix Real Space Refine. The model from Phenix Real Space refinement was taken and used in a final round of Rosetta FastRelax in torsion space to remove the various clashes that were introduced during Phenix refinement.

### ATPase Assays

Hsp104, _PARL_Skd3, and _PARL_Skd3 variants (0.25 µM monomer) were incubated with ATP (1 mM) (Innova Biosciences) at 37°C for 5 min in luciferase reactivation buffer (LRB; 25 mM HEPES-KOH [pH = 8.0], 150 mM KAOc, 10 mM MgAOc, 10 mM DTT). ATPase activity was assessed via inorganic phosphate release with a malachite green detection assay (Expedeon) and measured in Nunc 96 Well Optical plates on a Tecan Infinite M1000 plate reader. Background hydrolysis was measured at time zero and subtracted (Cupo and Shorter, 2020b; DeSantis et al., 2012). ATPase kinetics for _PARL_Skd3 was calculated using GraphPad Prism with a Michaelis-Menten least squares fit which was subsequently used to derive K_M_ and V_max_.

### Luciferase Disaggregation and Reactivation Assays

Firefly luciferase was aggregated by incubating luciferase (50 µM) in LRB (pH=7.4) with 8M urea at 30°C for 30 min. The denatured luciferase was then rapidly diluted 100-fold into ice-cold LRB, snap frozen, and stored at −80°C until use. Hsp104 was incubated with 50 nM aggregated firefly luciferase in the presence or absence of Hsc70 and Hdj2 (0.167 µM each) in LRB plus 5 mM ATP plus an ATP regeneration system (ARS; 1 mM creatine phosphate and 0.25 µM creatine kinase) at 37°C for 90 min (unless otherwise indicated). _PARL_Skd3 and variants (1 µM monomer, unless otherwise indicated) were incubated with 50 nM aggregated firefly luciferase in LRB plus 5 mM ATP plus ARS at 37°C for 90 min (unless otherwise indicated). Nucleotide-inhibitor assays for _PARL_Skd3 disaggregation activity were tested in the presence of ATP (Sigma), ATPγS (Roche), or ADP (MP Biomedicals) for 30 min at 37°C without ARS. IC_50_ curves for ADP and ATPγS were fitted using GraphPad Prism with a variable slope (four parameters) least squares fit. Recovered luminescence was monitored in Nunc 96 Well Optical plates using a Tecan Infinite M1000 plate reader (Cupo and Shorter, 2020b; DeSantis et al., 2012; Glover and Lindquist, 1998). Typically, Hsp104, Hsc70, and Hdj2 recovered ∼10% of native luciferase activity, whereas _PARL_Skd3 recovered ∼45% native luciferase activity (Cupo and Shorter, 2020b). Under our conditions, neither ADP nor ATPγS had an inhibitory effect on native luciferase (Figure 5).

### FITC-Casein Binding Assays

Fluorescence polarization was performed essentially as described previously (Rizo et al., 2019). _PARL_Skd3 was exchanged into 40 mM HEPES-KOH pH 8.0, 20 mM MgCl_2_, 150 mM KCl, 10% Glycerol (v/v), 2 mM β-mercaptoethanol. To assess FITC-casein binding, FITC-casein (60 nM, Sigma) was incubated with increasing concentrations (0-2.5µM hexameric) of _PARL_Skd3 with 2 mM of the indicated nucleotide for 10 min at 25°C. For the ATP condition, an ATP regeneration system (5 mM creatine phosphate and 0.125 µM creatine kinase) was included to maintain 2 mM ATP. Fluorescence polarization was measured (excitation 470 nm, emission 520 nm) using a Tecan Infinite M1000 plate reader. The binding isotherms were analyzed using Prism.

### Modeling Heterohexamer Ensemble Activity

The binomial distribution was used to model the activity of various heterohexamer ensembles (DeSantis et al., 2012; Werbeck et al., 2008):

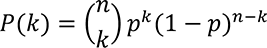

Where: P(k) is the probability of a hexamer containing k mutant subunits, n is total number of subunits (which for a hexamer, n=6), and p is the probability that a mutant subunit is incorporated. FRET subunit mixing experiments demonstrated that mutant and WT subunits have a similar probability of being incorporated into a hexamer (Figures S5D, 6A, B). Thus, p is calculated as the molar ratio of mutant and WT protein present:

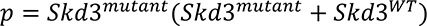

Therefore, for any specified concentration of mutant protein, the probability distribution of _PARL_Skd3 hexamers containing zero, one, two, three, four, five, or six mutant subunits can be derived (Figure 6A). Activity versus p plots (Figure 6B) can then be generated assuming each WT subunit makes an equal contribution to the total activity (one-sixth per subunit). Consequently, if subunits within the hexamer operate independently then activity should decline linearly upon incorporation of mutant subunits. Conversely, if subunit activity is coupled then the incorporation of a specific number of subunits will be sufficient to abolish activity. In our model, zero activity is assigned to hexamers that exceed the specific threshold number of mutant subunits. In this way, we generate activity versus p plots by assuming that 1 or more, 2 or more, 3 or more, 4 or more, or 5 or more mutant subunits are required to eliminate activity. This formula can be expressed as:

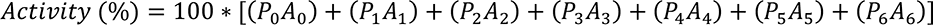

Where: P(k) is the probability of hexamer containing k mutant subunits derived above and A(k) is the relative assigned contribution to activity of a hexamer containing k mutant subunits.

### Alexa Fluor Labeling of _PARL_Skd3

For Förster resonance energy transfer (FRET) studies, we labeled separate pools of _PARL_Skd3 with Alexa-Fluor 488 (Alexa488, ThermoFisher Scientific CAT# A20000) as the FRET donor and Alexa-Fluor 594 (Alexa594, ThermoFisher Scientific CAT# A20004) as the FRET acceptor. In brief, _PARL_Skd3 (WT and mutants) was extensively exchanged into labelling buffer (LB; 50 mM HEPES-KOH [pH = 8.0], 150 mM KCl, 20 mM MgCl_2_, 10% glycerol, 10mM BME) at room temperature using Micro Bio-Spin 6 columns (Bio-Rad CAT# 7326200). _PARL_Skd3 concentration was measured via A280 and the molar extinction coefficient and _PARL_Skd3 concentration was adjusted to 30 μM. The primary amine (R-NH2) reactive dye, Alexa-Fluor 488 NHS Ester (Succinimidyl Ester) (Alexa488) or Alexa-Fluor 594 NHS Ester (Succinimidyl Ester) (Alexa594), was then added to _PARL_Skd3 samples to achieve a 10-fold molar excess over _PARL_Skd3. Samples were then incubated at 25°C in the dark. After 75min, the labeling reaction was quenched by rapidly and extensively exchanging into labelling buffer + 10 mM DTT. To ensure that all unreacted dye is removed, the buffer exchange step was repeated at least twice. _PARL_Skd3 concentration and labeling efficiency were determined by UV/Vis spectrometry according to the manufacturer’s instructions (Invitrogen). Typically, we achieved ∼50% labelling efficiency.

### Fluorescence Resonance Energy Transfer and Subunit Mixing

We employed Förster resonance energy transfer (FRET) to measure subunit mixing (Figure S5D, 6A, and 6B). Donor (Alexa-Fluor 488) labeled _PARL_Skd3 (Alexa488-_PARL_Skd3), Acceptor (Alexa-Fluor 594) labeled _PARL_Skd3 (Alexa594-_PARL_Skd3), or free dye were mixed in equal stoichiometric parts to a final total dye concentration of 1μM in labelling buffer with ATP (5 mM). Because these dyes function as a FRET pair with a Förster radius of 60 Å, primary amine labelling in _PARL_Skd3 is expected to yield intermolecular FRET once mixed oligomers are formed. Given the R_0_ value of 60Å for the Alexa488-Alexa594 FRET pair, it is possible to observe both intrahexameric FRET (e.g. solvent exposed residues K538 from P3 and K658 from P4 are 8.5 Å apart) and interhexameric FRET within a dodecamer (closest two lysines are K134 to K265 at 22.4 Å apart, but it is unclear if they are solvent exposed). However, the hexameric state is the predominant species in the absence of substrate and thus our FRET assay likely reports on the hexameric state of _PARL_Skd3 rather than the dodecamer (Figure 1B). A similar strategy has been employed to demonstrate the formation of mixed hexamers by bacterial ClpB (Werbeck et al., 2008), Hsp104 (DeSantis et al., 2012), and MCM helicase, another AAA+ ATPase, from *Sulfolobus solfataricus* (McGeoch et al., 2005; Moreau et al., 2007). Prior to any measurements, samples were allowed to equilibrate for 10 min at room temperature. Mixed and equilibrated samples were excited at the donor excitation wavelength of 480nm. Donor fluorescence was measured at 519nm with a bandwidth of 5nm. Acceptor fluorescence was measured at 630nm with a bandwidth of 5nm. Apparent FRET efficiency (Figure S5D, S6C, D) was calculated from Alexa488-_PARL_Skd3 emission (488nm) and Alexa594-_PARL_Skd3 emission (519nm) as F_a_/(F_d_+F_a_), where F_d_ is the measured Alexa488-_PARL_Skd3 (donor) fluorescence and F_a_ is the Alexa594-_PARL_Skd3 (acceptor) fluorescence in the presence of Alexa488-Skd3.

### Data Availability

_PARL_Skd3:casein:ATPγS cryo-EM maps and atomic coordinates have been deposited in the EMDB and PDB with accession codes EMDB-26121 (State 1), EMDB-26122 (State 1 filtered), PDB 7TTR (State 1, AAA+ only), and PDB 7TTS (State 1, Full Model).

## Acknowledgements

We thank JiaBei Lin, Linamarie Miller, Charlotte Fare, Katie Copley, and Zarin Tabassum for critiques. This work was funded by the Blavatnik Family Foundation Fellowship (R.R.C.), The G Harold and Leila Y Mathers Foundation (J.S.), and NIH grants T32GM008275 (R.R.C.), F31AG060672 (R.R.C.), R01GM099836 (J.S.), R21AG061784 (J.S.), and R01GM138690 (D.R.S). J.S. is a consultant for Dewpoint Therapeutics, Vivid Sciences, Neumora, Korro Bio, and ADRx.

---

**Figure S1.**
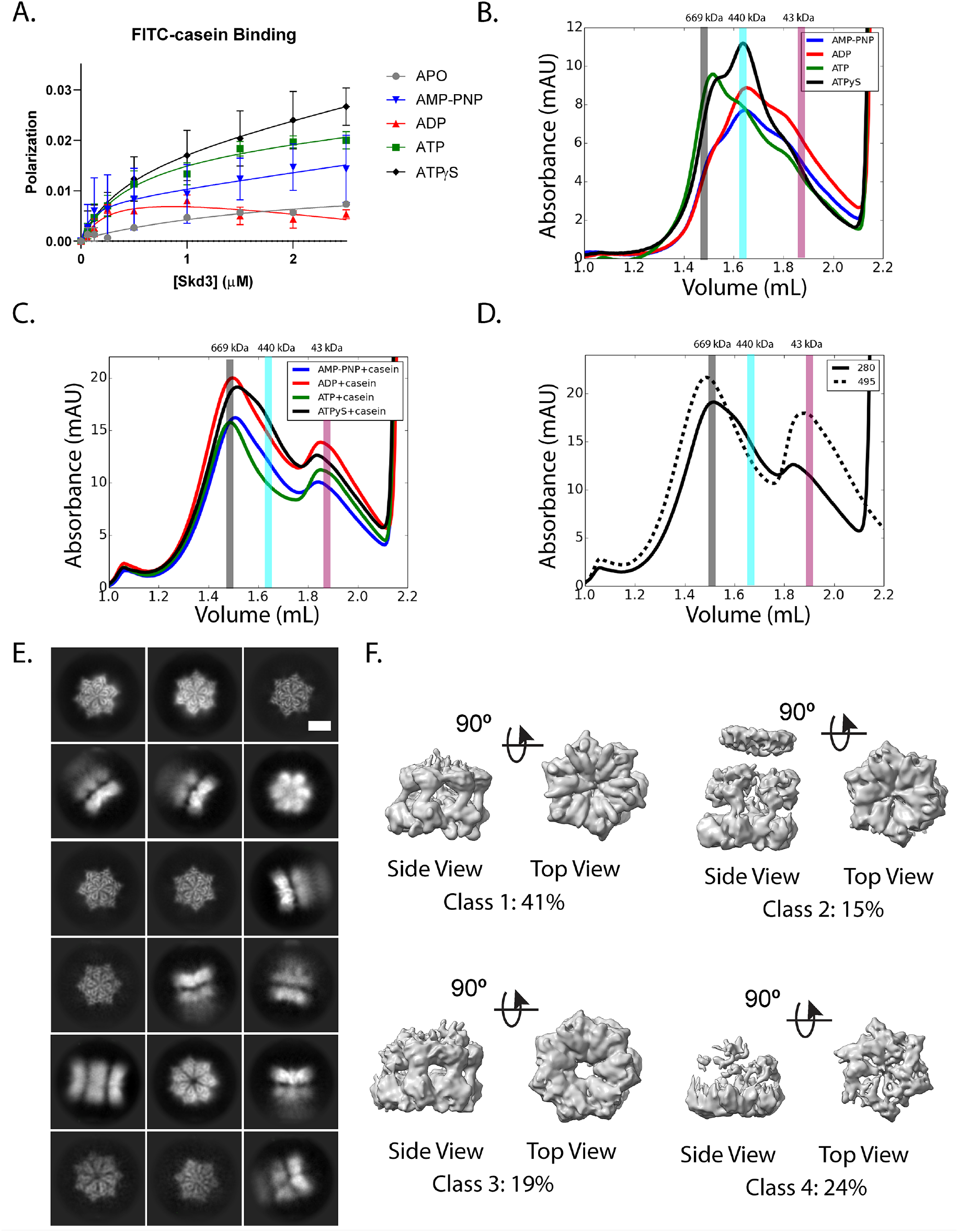
Structure of _PARL_Skd3. **(A)** FITC-casein binding analysis, measured by fluorescence polarization in the presence of no nucleotide (APO; grey), AMP-PNP (blue), ADP (red), ATP (green), and ATPγS (black). Values represent means ± SEM (n=3). **(B, C)** SEC-trace of _PARL_Skd3 with different nucleotides without casein **(B)** and with casein **(C)** including AMP-PNP (blue), ADP (red), ATP (green), and ATPγS (black). The three vertical bars represent different molecular weight standards thyroglobulin (669 kDa), ferritin (440 kDa), and ovalbumin (43 kDa) that approximately represent Skd3 dodecamers (grey), hexamers (cyan), or monomers (magenta). **(D)** SEC-trace of _PARL_Skd3:casein:ATPγS complex with both 280nm UV absorbance (solid) and 495nm UV absorbance to detect FITC-casein (dashed) shown. Vertical bars indicate molecular-weight standards as in Figure S1B. **(E)** Representative 2D class averages from the full dataset. Scale bar, 100Å. **(F)** 3D classification results from the total combined dataset: hexamer, Class 1 (top, left); dodecamer, Class 2 (top, right); heptamer, Class 3 (bottom, left); and other, Class 4 (bottom, right). Related to Figure 1.

**Figure S2.**
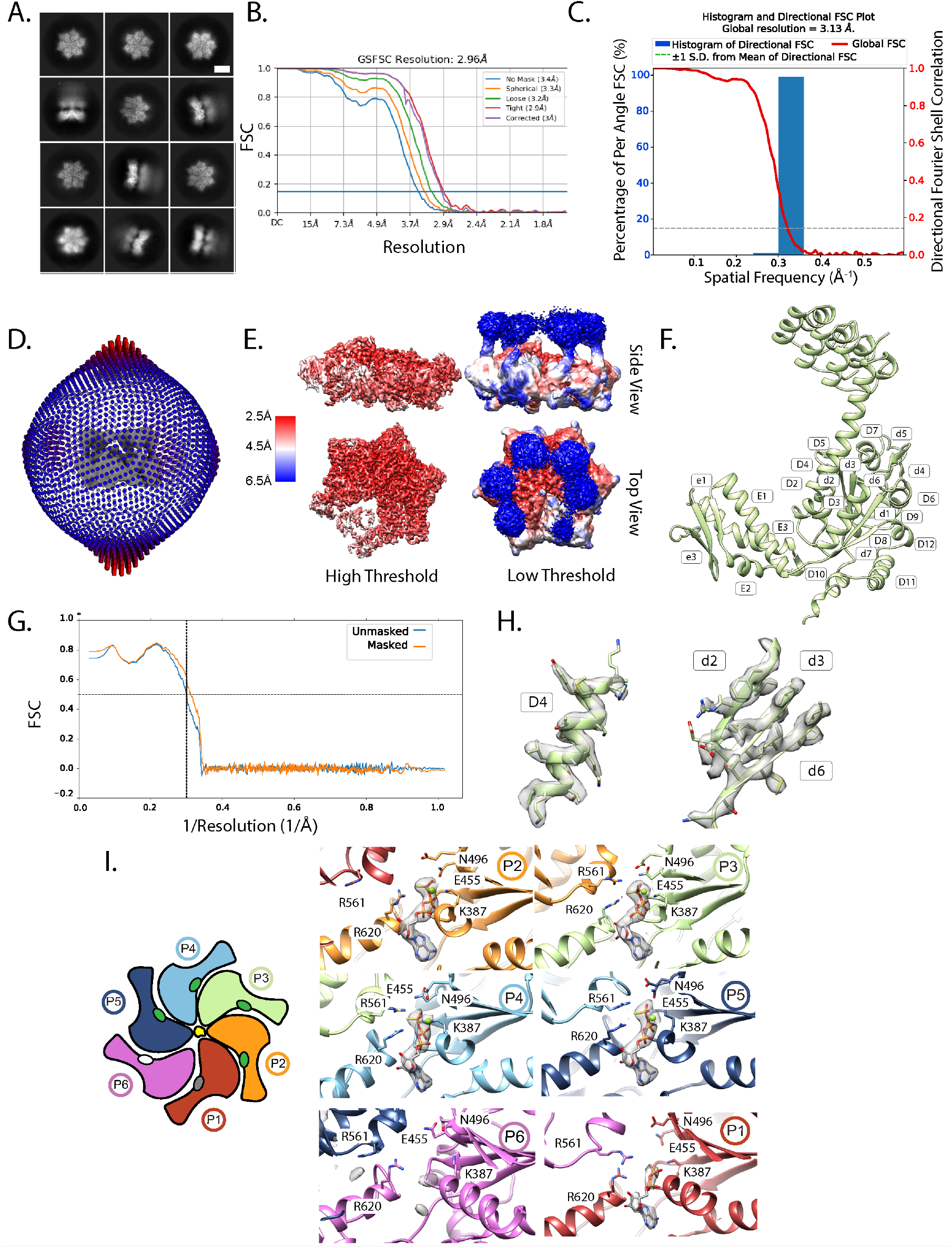
Structure and nucleotide-binding pockets of _PARL_Skd3. **(A)** Representative 2D class averages from the hexamer class. Scale bar, 100Å. **(B)** Gold standard FSC-curves for the hexamer class refinement. **(C)** Histogram and directional FSC plot for the hexamer class. **(D)** Particle distribution map of the dodecamer class. **(E)** Local resolution map of the hexamer map and two different threshold high threshold (left) and low threshold (right). **(F)** _PARL_Skd3 structure labeled for reference with NBD helices and strands indicated for the large subdomain (D) or small subdomain (E) as for NBD2 of bacterial ClpB (PDB: 1QVR) (Lee et al., 2003). **(G)** Map vs. Model FSC for both unmasked (blue) and masked (orange) for the hexamer model. **(H)** Map plus model of alpha helix, D4, and beta-sheets d2, d3, and d6. **(I)** Schematic of overall NBD structure (top view) with circles representing ATP (green), ADP (grey), or APO (white) in the nucleotide-binding pocket (left). Map and model of the nucleotide-binding pocket with residues involved in ATP hydrolysis are shown and labeled including Arg Finger (R561), sensor-1 (N496), sensor-2 (R620), Walker A (K387), and Walker B (E455) (right). Related to Figure 1 and 2.

**Figure S3.**
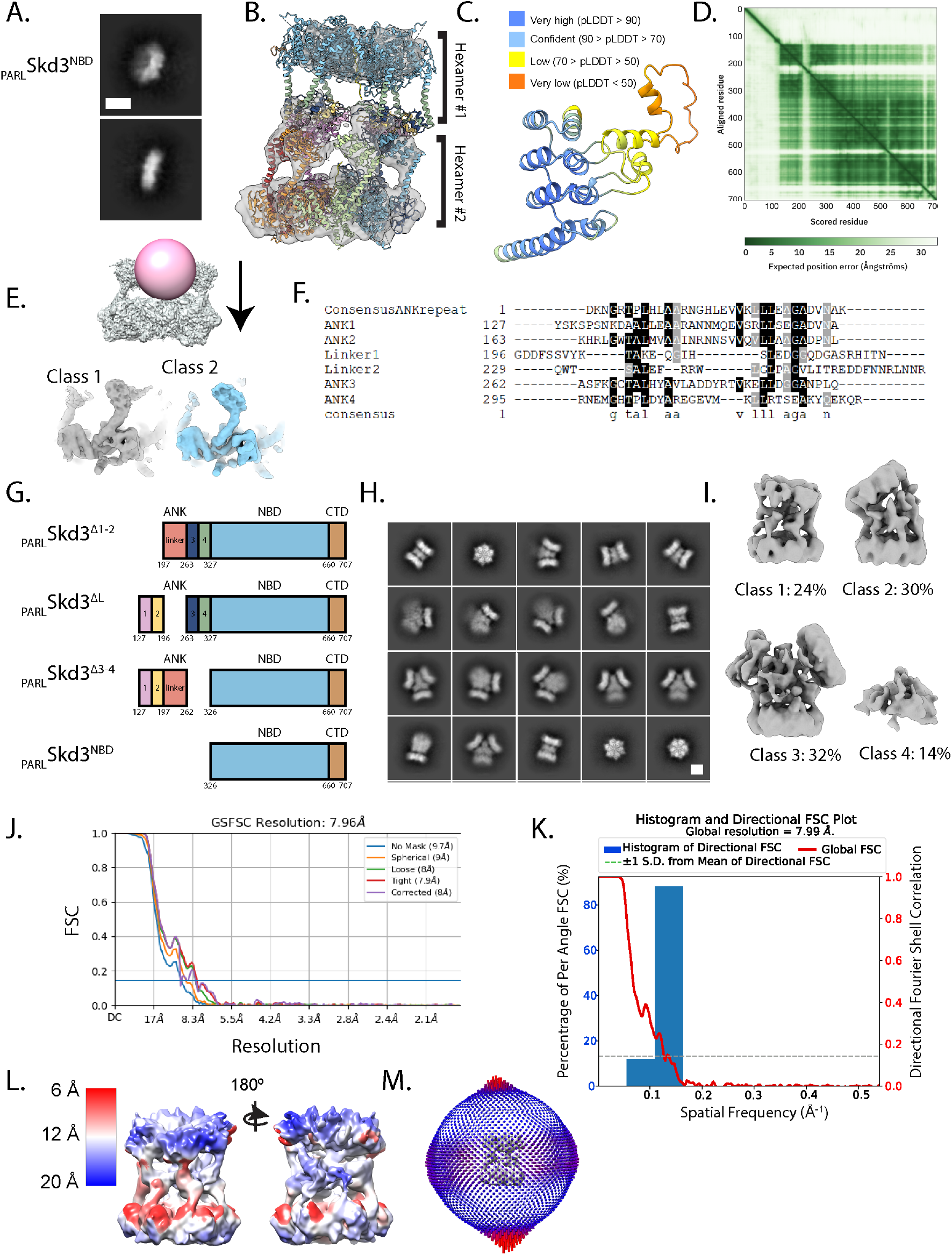
Structural refinement of the ANK. **(A)** Representative 2D class averages from _PARL_Skd3^NBD^. Scale bar, 100Å. **(B)** Dodecamer map of _PARL_Skd3 from 3D classification with representative model colored by individual domains docked in the top and bottom of the hexamer. **(C)** Model prediction from AlphaFold of ANK colored by pLDDDT score. **(D)** Plot of the predicted aligned error of the AlphaFold prediction of full length Skd3. **(E)** Mask (pink) and map (grey) used in focus classification in cisTEM of the single ankyrin domain on the hexamer class (left) with results of two representative classes (right). **(F)** Alignment of the four ankyrin repeats and the linker region of *H. sapiens* Skd3 to the consensus ankyrin repeat from Mosavi, *et. al.* (Mosavi et al., 2002). Alignments were constructed using Clustal Omega. Linker region was aligned to consensus sequence manually. Bottom row shows consensus sequence of alignment. **(G)** Domain architecture maps of the different ANK deletion mutations. **(H)** Representative 2D class averages from the _PARL_Skd3^Δ1-2^ dataset. Scale bar, 100Å. **(I)** 3D classification results from the _PARL_Skd3^Δ1-2^ dataset: dodecamer, Class 1 (left); bent dodecamer, Class 2 (middle left); trimer, Class 3 (middle right); and other, Class 4 (right). **(J)** Gold standard FSC-curves for the final dodecamer class refinement. **(K)** Histogram and directional FSC plot for the dodecamer class. **(L)** Local resolution map of the dodecamer map. **(M)** Particle distribution map of the dodecamer class. Related to Figure 3.

**Figure S4.**
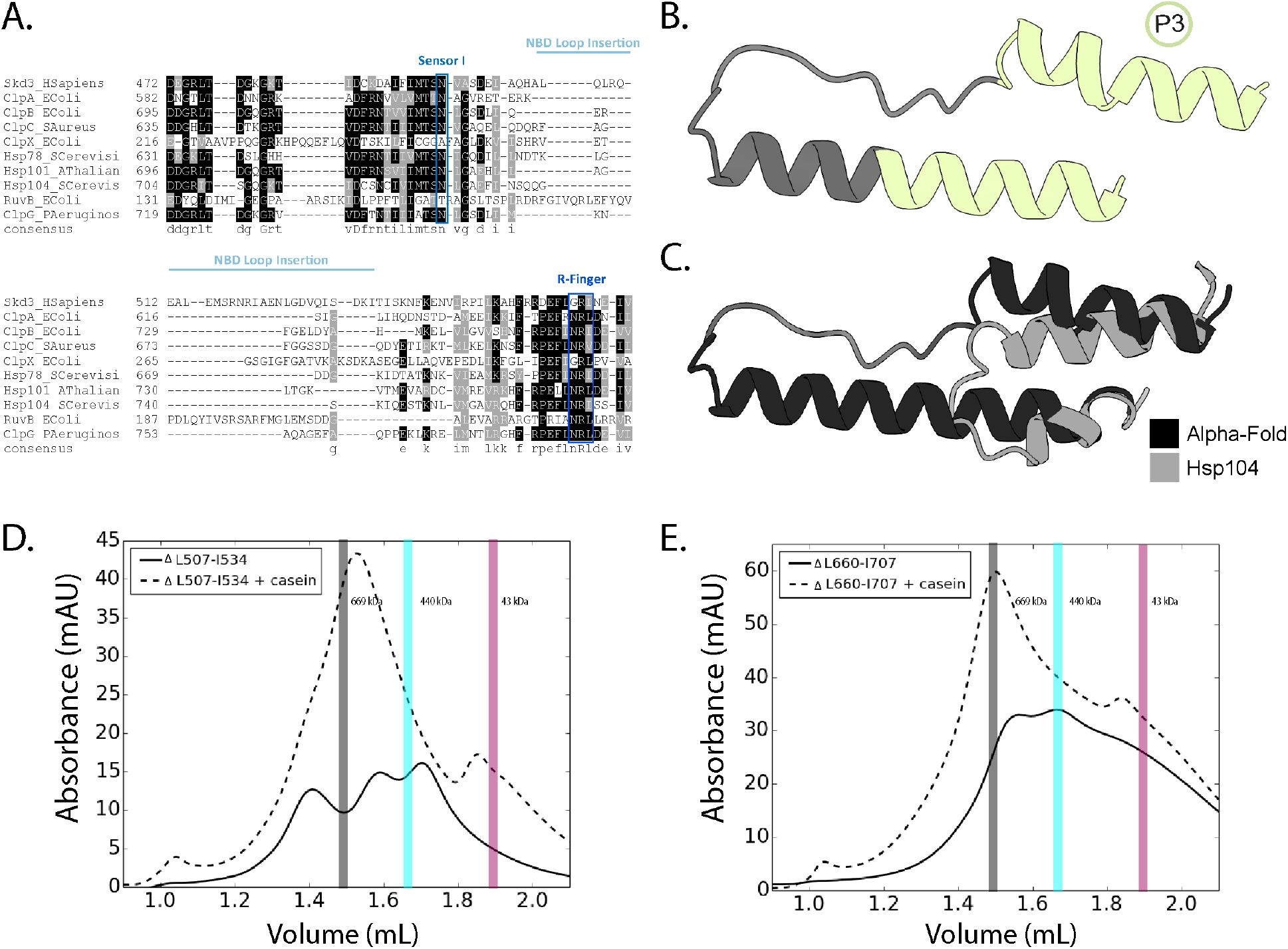
Structural features of the NBD insertion and CTD. **(A)** Alignment of a select region of the NBD from *H. sapiens* Skd3, NBD2 from *E. coli* ClpA, NBD2 from *E. coli* ClpB, NBD from *A. aureus* ClpC, NBD from *E. coli* ClpX, NBD2 from *S. cerevisiae* Hsp78, NBD2 from *A. thaliana* Hsp101, NBD2 from *S. cerevisiae* Hsp104, NBD from *E. coli* RuvB, and NBD2 from *P. aeruginosa* ClpG. Alignments were constructed using Clustal Omega. Bottom row shows consensus sequence of alignment. Highlighted in blue are the sensor-1 and Arg-finger motifs. Light blue highlights the insertion from L507-I534 in the Skd3 NBD. **(B)** Alpha-fold model prediction (grey) of the NBD insertion alone (residues 449-515 to 535-552 are shown). The residues that were successfully built in de novo are represented in green on the model of protomer 3. **(C)** The Alpha-fold model prediction (black) overlayed with the Hsp104 model (grey, PDB: 5VJH). **(D)** SEC of _PARL_Skd3^ΔL507-I534^ plus ATPSγS with casein (dashed) and without casein (solid). The three vertical bars represent different molecular weight standards thyroglobulin (669 kDa), ferritin (440 kDa), and ovalbumin (43 kDa) that approximately represent Skd3 dodecamers (grey), hexamers (cyan), or monomers (magenta). **(E)** SEC of _PARL_Skd3^ΔL660-I707^ plus ATPSγS with casein (dashed) and without casein (solid). The three vertical bars represent different molecular weight standards thyroglobulin (669 kDa), ferritin (440 kDa), and ovalbumin (43 kDa) that approximately represent Skd3 dodecamers (grey), hexamers (cyan), or monomers (magenta). Related to Figure 4.

**Figure S5.**
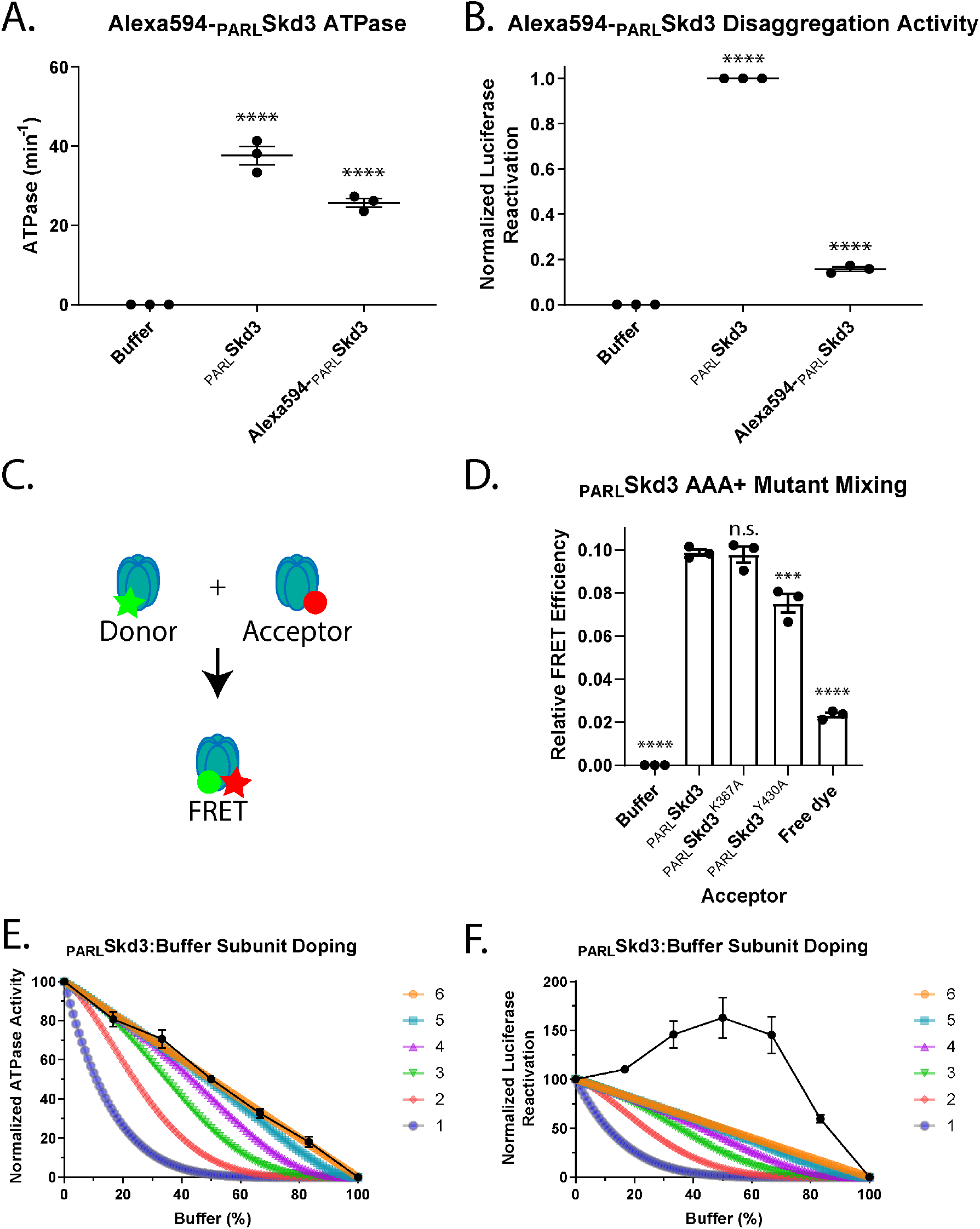
Skd3 is a subglobally cooperative protein disaggregase. **(A)** ATPase activity of _PARL_Skd3 and Alexa594-_PARL_Skd3. ATPase activity was compared to buffer using one-way ANOVA and a Dunnett’s multiple comparisons test (N = 3, individual data points shown as dots, bars show mean ± SEM, ****p<0.0001). **(B)** Luciferase disaggregase activity of _PARL_Skd3 and Alexa594-_PARL_Skd3. Luciferase activity was buffer subtracted and normalized to _PARL_Skd3. Disaggregase activity was compared to buffer using one-way ANOVA and a Dunnett’s multiple comparisons test (N = 3, individual data points shown as dots, bars show mean ± SEM, ****p<0.0001). **(C)** Schematic of subunit mixing assayed by FRET. Separate pools of _PARL_Skd3 were labeled with Alexa-Fluor 488 (Alexa488) to serve as a donor and Alexa-Fluor 594 (Alexa594) to serve as an acceptor. In mixed hexamers, the donor (Alexa488) and acceptor (Alexa594) labels come into close enough proximity to elicit FRET. **(D)** FRET efficiency after mixing Alexa488-_PARL_Skd3 with buffer, Alexa594-_PARL_Skd3, Alexa594-_PARL_Skd3^K387A^, Alexa594-_PARL_Skd3^E455Q^, or Alexa594-_PARL_Skd3^Y430A^ for 10 min in the presence of ATP (5 mM) at a 1:1 molar ratio with a final labelled _PARL_Skd3 concentration of 1μM. As a negative control the FRET efficiency of mixing unreacted Alexa488 dye with unreacted Alexa594 dye is also shown. Relative FRET efficiency was compared to WT _PARL_Skd3 using one-way ANOVA and a Dunnett’s multiple comparisons test (N = 3, individual data points shown as dots, bars show mean ± SEM, ****p<0.0001). **(E)** ATPase activity of _PARL_Skd3 mixed with various ratios of buffer. ATPase activity was buffer subtracted and normalized to _PARL_Skd3 (N = 3, data shown as black dots with mean ± SEM). **(F)** Luciferase disaggregase activity of _PARL_Skd3 mixed with various ratios of buffer. Disaggregase activity was buffer subtracted and normalized to _PARL_Skd3 (N = 3, data shown as black dots with mean ± SEM). Related to Figure 6.

**Figure S6.**
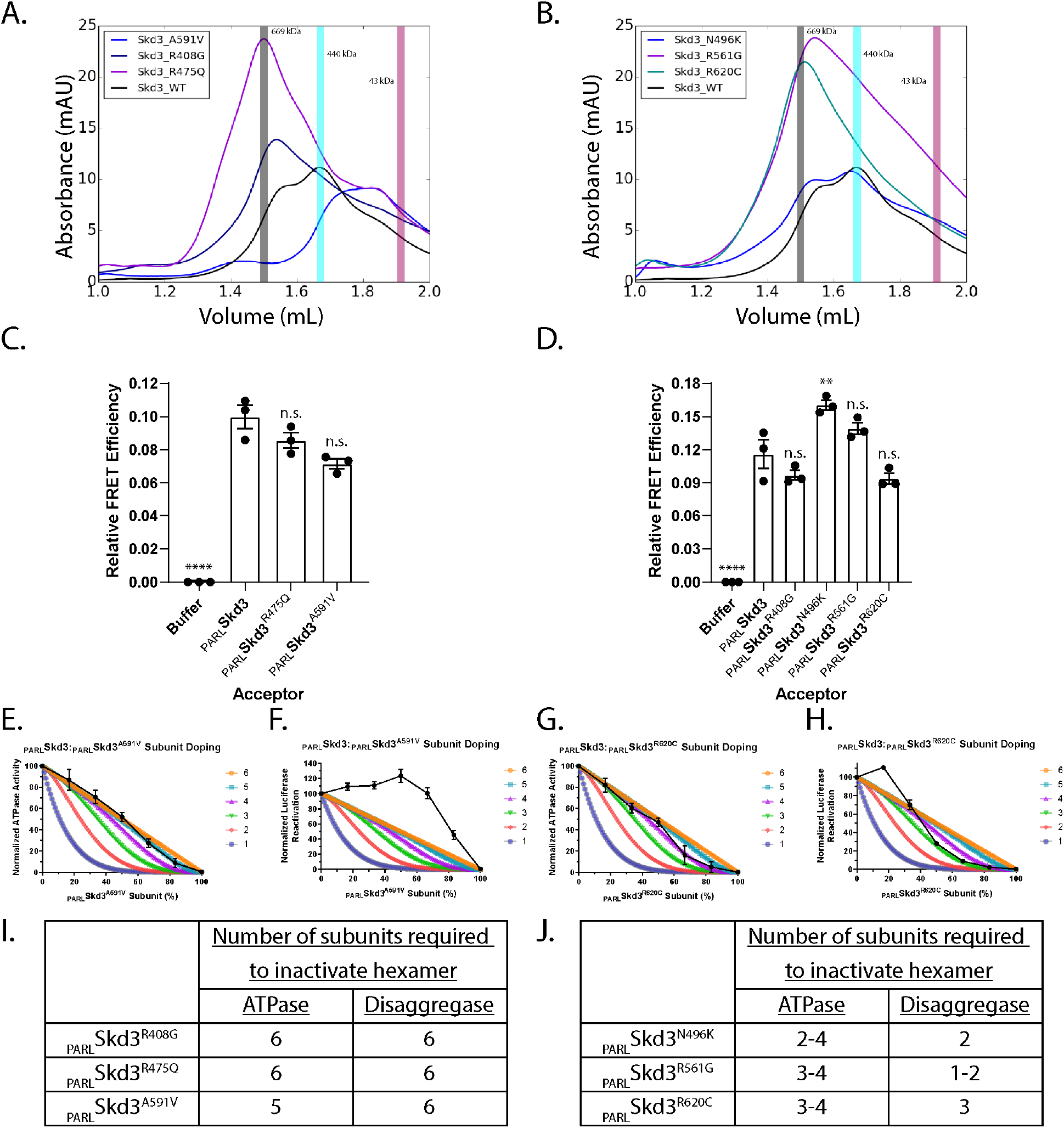
SCN-linked subunits inhibit _PARL_Skd3 activity more severely than MGCA7-linked _PARL_Skd3 subunits. **(A)** EC of MGCA7-linked _PARL_Skd3 variants. The three vertical bars represent different molecular weight standards thyroglobulin (669 kDa), ferritin (440 kDa), and ovalbumin (43 kDa) that approximately represent Skd3 dodecamers (grey), hexamers (cyan), or monomers (magenta). **(B)** SEC of SCN-linked _PARL_Skd3 variants. The three vertical bars represent different molecular weight standards thyroglobulin (669 kDa), ferritin (440 kDa), and ovalbumin (43 kDa) that approximately represent Skd3 dodecamers (grey), hexamers (cyan), or monomers (magenta). **(C)** FRET efficiency after mixing Alexa488-_PARL_Skd3 with buffer, Alexa594-_PARL_Skd3, Alexa594-_PARL_Skd3^A591V^, or Alexa594-_PARL_Skd3^R475Q^ for 10 min in the presence of ATP (5 mM) at a 1:1 molar ratio with a final labeled _PARL_Skd3 concentration of 1μM. Relative FRET efficiency was compared to WT _PARL_Skd3 using one-way ANOVA and a Dunnett’s multiple comparisons test (N = 3, individual data points shown as dots, bars show mean ± SEM, ****p<0.0001). **(D)** FRET efficiency after mixing Alexa488-_PARL_Skd3 with buffer, Alexa594-_PARL_Skd3, Alexa594-_PARL_Skd3^R408G^, Alexa594-_PARL_Skd3^N496K^, Alexa594- _PARL_Skd3^R561G^, or Alexa594-_PARL_Skd3^R620C^ for 10 min in the presence of ATP (5 mM) at a 1:1 molar ratio with a final labelled _PARL_Skd3 concentration of 1μM. Relative FRET efficiency was compared to WT _PARL_Skd3 using one-way ANOVA and a Dunnett’s multiple comparisons test (N = 3, individual data points shown as dots, bars show mean ± SEM, **p<0.01, ****p<0.0001). **(E-H)** ATPase activity of _PARL_Skd3 was mixed with various ratios of _PARL_Skd3^A591V^ (E) or _PARL_Skd3^R620C^ (G). ATPase activity was buffer subtracted and normalized to _PARL_Skd3 (N = 3, data shown as black dots with mean ± SEM). Luciferase disaggregase activity of _PARL_Skd3 mixed with various ratios of _PARL_Skd3^A591V^ (F) or _PARL_Skd3^R620C^ (H). Disaggregase activity was buffer subtracted and normalized to _PARL_Skd3 (N = 3, data shown as black dots with mean ± SEM). **(I)** Table summarizing the effect of MGCA7-linked subunits on ATPase activity and luciferase disaggregase activity. **(J)** Table summarizing the effect of SCN-linked subunits on ATPase activity and luciferase disaggregase activity. Related to Figure 7.

**Table S1.** Cryo-EM data collection, refinement and validation statistics of _PARL_Skd3 structures.

|  | PARLSkd3<br>(Class 1,AAA+ only)<br>EMDB 26121<br>PDB 7TTR | PARLSkd3<br>(Class 1, AAA+ & ANK)<br>EMDB 26122<br>PDB 7TTS | PARLSkd3<br>(Class 2) | PARLSkd3<sup>Δ1-2</sup><br>(Class 1) |
| --- | --- | --- | --- | --- |
| **Data collection and processing** |  |  |  |  |
| Microscope and camera | Titan Krios, K3 |  |  | Glacios, K2 |
| Magnification | 105,000 |  |  | 45,000 |
| Voltage (kV) | 300 |  |  | 200 |
| Data acquisition software | Serial EM |  |  | Serial EM |
| Exposure navigation | Image Shift |  |  | Image Shift |
| Electron exposure (e<sup>-</sup>/Å<sup>2</sup>) | 68 |  |  | 55.8 |
| Final particle images (no.) | 358,000 |  | 130,000 | 165,354 |
| Map resolution (Å) | 2.96 | ~6 (filtered) | 7.2 | 7.9 |
| FSC threshold | 0.143 |  |  |  |
| Map resolution range (Å) | 2.5-6.5 | 6 | 6-10 | 6-20 |
| **Refinement** |  |  |  |  |
| Model resolution (Å) | 2.9 | 2.9 | - | - |
| FSC threshold | .143 | .143 | - | - |
| Map sharpening *B* factor (Å<sup>2</sup>) | -119 | - | -497 | -610 |
| **Model composition** |  |  |  |  |
| Non-hydrogen atoms | 15,699 | 18,796 | - | - |
| Protein residues | 1926 | 2700 | - | - |
| Ligands | 9 | 9 | - | - |
| **B factors (Å<sup>2</sup>)** |  |  | - | - |
| Protein | 158.16 | 178.33 | - | - |
| Ligand | 54.10 | 54.10 | - | - |
| **R.m.s deviations** |  |  |  |  |
| Bond lengths (Å) | .011 | .010 | - | - |
| Bond angles (°) | 1.249 | 1.222 | - | - |
| **Validation** |  |  |  |  |
| MolProbity score | 1.67 | 1.53 | - | - |
| Clashscore | 6 |  | - | - |
| Poor rotamers (%) | .30 | .30 | - | - |

**Movie S1. Structure of _PARL_Skd3 bound to casein in the presence of ATPγS.**

**Movie S2. Structure of _PARL_Skd3^Δ1,2^ bound to casein in the presence of ATPγS.**

